# The Past, Present and Future of Elephant Landscapes in Asia

**DOI:** 10.1101/2020.04.28.066548

**Authors:** Shermin de Silva, Tiffany Wu, Philip Nyhus, Alison Thieme, Ashley Weaver, Josiah Johnson, Jamie Wadey, Alexander Mossbrucker, Thinh Vu, Thy Neang, Becky Shu Chen, Melissa Songer, Peter Leimgruber

## Abstract

Habitat loss drives species’ declines worldwide, but is seldom quantified over centennial timescales. We constructed ecological niche models for Asian elephants based on land-use change between 850-2015, and predictions under six different climate/socioeconomic scenarios from 2015-2099. We find that over 64% of suitable natural habitat across diverse ecosystems was lost over the past three centuries. Average patch size dropped 83% from approximately 99,000 km^2^ to 16,000 km^2^ and the area occupied by the largest patch decreased 83% from ~ 4 million km^2^ (45% of area) to 54,000 km^2^ (~7.5% of area). Over half of current elephant range appears unsuitable. Habitat availability is predicted to decline further this century across all scenarios. The most severe losses occur under RCP3.4-SSP4, representing mid-range emissions but high regional inequities. We conclude that climate change mitigation measures must include policies to ensure inter-regional socioeconomic equity to safeguard landscapes for elephants, humans, and other species.

## Introduction

Habitat loss and degradation are leading drivers of terrestrial biodiversity loss worldwide ^1–3^. An estimated three quarters of the Earth’s land surface has been significantly altered by human activities ^4^. Historic reasons include conversion for cultivation and settlement, reflecting both local and global socioeconomic drivers of land-use and land-cover (LULC) change ^5,6^. Climate change is an additional contributor to species declines within the past century ^4,6^. As a result of these anthropogenic changes to climate and land-use, global forest extent has been reduced by 32% relative to the pre-industrial period and ecological communities are estimated to have lost over 20% of their biodiversity ^4,6^. Yet it is not a given that efforts to address climate change will also conserve species and habitats because these are distinct concerns ^7^. Moreover, climate change and LULC feed back on each other via economic systems as well as natural processes such as carbon cycling ^5^. These social-ecological systems impose inherent trade-offs on efforts to mitigate climate change and conserve biodiversity ^7,8^. Indeed, simultaneously addressing both goals may be among the greatest and most urgent challenges to developing sustainable societies ^6^.

Human activities can alter landscapes quickly, but long-term legacies may not be immediately evident. Human-induced changes already restrict the ranges of many terrestrial mammal species ^9,10^ but historical records on population abundance and distribution are often limited for many taxa, complicating efforts to assess human impacts over longer periods. Though LULC trends in recent decades may be inferred from satellite imagery ^2,11^, these fail to capture processes operating over centuries. Nevertheless, longer historical perspectives are necessary to appreciate the true magnitude of changes to threatened ecosystems. For example, historical studies have influenced conservation policies related to remnant prairies in Oregon, wetlands in Iowa, and forests in Germany, at times challenging traditional management practices ^12^. However, a review of more than 200 historical ecology studies found that fewer than 12% contained recommendations that “explicitly addressed ongoing or projected climate change” ^12^. Moreover, although many studies attempt to define global priority regions for conservation ^7,13^, they do not account for future changes in climate, socioeconomic conditions, and land-use. There is a need for better integration of historical perspectives and present-day assessments with forward-looking strategies and policies.

Land-use practices and climate change will continue to alter ecosystem structure and in the future, resulting in changes to the availability of habitat for many species ^1^. Wide-ranging species cannot be conserved solely within existing or planned protected area networks ^1^. The survival of these species will depend not only on reducing proximate threats from illegal hunting and sufficient protection of extant range, but on preparing for possible range shifts and associated challenges under scenarios of global change. Typically, this is achieved through ecological niche modelling (c.f. species distribution modelling) in which present-day species occurrence data, together with environmental covariates, are used to infer possible occurrence or suitable habitat at a different area or time ^14–16^. Often, climatic variables form the basis of predictive studies^1^ but LULC features more directly constrain habitat availability. Moreover, social and economic systems can be expected to evolve along with physical changes in the climate and biosphere ^17^. Therefore, biodiversity conservation and land-use planning efforts must take into consideration not only possible global emissions outcomes, but also the corresponding LULC and socioeconomic pathways taken to reach those outcomes.

Asia, Earth’s most densely populated yet biodiverse continent, contains several hotspots of species decline and threatened megafauna ^9^. We model historic range in suitable habitat over the past 300 years and possible shifts over the next 100 years for a widely-distributed endangered mammal, the Asian elephant (*Elephas maximus*). Elephants are ecosystem engineers that uniquely influence the structure of natural ecosystems ^18^ and also exemplify mutual challenges for people and wildlife at frontiers of land-use change that manifest as human-elephant conflict ^19^. Occupying ecosystems ranging from rainforests to grasslands, this habitat generalist acts as a surrogate for diverse ecoregions rather than particular biomes. This range is thought to represent only a subset of past distributions, reflecting long-term contraction (Figure S1) ^20^. However, published depictions of past range are based on limited historical and anecdotal accounts whereas elephants were probably never distributed uniformly over the area.

We base our predictions on the Land-Use Harmonization 2 (henceforth LUH2) data products ^21^. The LUH2 datasets provide historical reconstructions of land-use and management between 850-2015, at annual increments. They also contain modelled scenarios from 2015-2100 representing combinations of a subset of Representative Concentration Pathways (RCPs)^22^ and Shared Socioeconomic Pathways (SSPs)^17^. These datasets therefore link historic and future land-use patterns with a consistent set of variables, and pair the socioeconomic narratives with possible emissions outcomes (Table 1; supplementary text section 1). We used a subset of LUH2 variables to model changes in habitat for elephants over multiple centuries (Table S1).

**Table 1.** Scenario summaries. Representative Concentration Pathway (RCP) number denotes the radiative forcing target in W/m^2^ achieved by a given pathway at stabilization by the year 2100. The Shared Socioeconomic Pathway (SSP) number denotes the specific narrative storyline that describes the socioeconomic processes operating in the modelled hypothetical societies. Here we describe features that are salient to land-use. See references ^17,21,39^ as well as in-table references and supplementary text section 1 for additional details.

| Scenario | Description |
| --- | --- |
| **RCP 2.6-**<br>**SSP 1:** | Warming stays below $2^\circ\text{C}$ by the end of the century, with climate policy implemented through carbon pricing. This sustainable development paradigm is characterized by the adoption of sustainable technologies including carbon capture and storage together with biofuel production and reforestation. The extent of protected areas globally achieves the 17% Aichi target. Global population levels off at mid-century with concurrent adoption of less resource-intensive and environmentally responsible lifestyles such as the consumption of less meat (thus requiring less pasture). Likewise, improvements in agricultural productivity and irrigation efficiency reduce the agricultural footprint. Economic development is high, along with strong governance and institutions at both national and international levels. |
| **RCP 3.4-**<br>**SSP 4:** | Mid-range radiative forcing target in a world of deep regional inequalities both within and among countries. High-income countries enjoy technological progress including agricultural innovations improving efficiency, concurrent with greater demands for energy and food. The high carbon price results in $\sim 1200$ million ha of land being allocated to the production of bioenergy, driving an 80% increase in the extent of cropland between 2010-2100. Climate and environmental policies in middle/high-income regions yield increases in forest cover. However, low-income countries stagnate. Consumption is driven by population growth, while agricultural productivity remains low and there is a heavy reliance on traditional biofuels, resulting in higher rates of deforestation. |
| **RCP 4.5-**<br>**SSP 2:** | The RCP target is achieved by shifts toward more sustainable energy sources, food systems with reduced carbon footprint, and expansion of forests following an initial decline. SSP 2 is an intermediate between SSP 1 and SSP 3, where population growth slows and levels off and most countries are politically stable but development progress is uneven within and among countries. |
| **RCP 6-**<br>**SSP 4:** | An alternative to RCP3.4-SSP4 based on the same framework but assuming less stringent climate change mitigation policies. Crop and pasture land increase by 14% and 9% respectively by 2100, along with a 3% increase in forest cover globally. The latter is largely situated in middle- and high-income countries. |
| **RCP 7-**<br>**SSP 3:** | SSP3 represents weak global institutions, nationalistic political and economic policies reduce international cooperation. Wealthier countries experience less population growth, others experience higher growth. Countries strive to meet their food and energy needs domestically. Slow technological change in the energy sector with continued reliance on fossil fuels. Land-use allocations are made through profit/productivity-maximizing decisions with unprotected and uncultivated areas more likely to be converted to other uses. |
| **RCP 8.5-**<br>**SSP 5:** | A fossil fuel-driven world with rapid economic development accompanied by investments in human capital, promoting more equitable distribution of resources, stronger institutions and slower population growth. |

Our aims are to: (1) quantify historic trends in suitable natural habitat and fragmentation over the 13 range countries throughout Asia; (2) characterize the suitability of present-day range and identify other areas containing potential habitat; and (3) predict how the extent and fragmentation of natural habitat may change under the six different RCP-SSP future scenarios. Under the simple assumption that climate trends are the critical determinant of habitat loss for this species, one would expect a decrease in suitable habitat with increasing RCP values. If socioeconomic considerations have more influence on land-use trends, there would be no such pattern. We highlight areas of concern and opportunity among the different regions, alongside the effect of climatic vs. socioeconomic pathways on elephant landscapes.

## Results

### Historical change

Primary forest was less important for the model than elevation, forested and non-forested secondary lands, croplands, pastures and wood harvest activities (Table S2). The amount ‘suitable’ natural habitat was relatively stable until the 1700s but afterwards declined by approximately 64% (over 38% within current range; Figures 1 & S2, Table 2). Average patch size fell by 83%, from 99,000 km^2^ to 16,000 km^2^ and amount of area occupied by the largest patch (LPI) also decreased 83% from 4 million km^2^ (~45% of total area) to a little over 54,000 km^2^ (~7.5% of total area). The landscape contagion index nearly doubled (Figure 1). Mainland China, India, Bangladesh, Thailand, Vietnam and Sumatra each lost more than half their suitable range (Table 2). China lost 94% and India lost 86%. Bhutan, Nepal and Sri Lanka lost more suitable habitat inside than outside current elephant range (Table 2). Trends in Lao PDR and Malaysia showed a net gain, but not necessarily in areas within the current range (Table 2 and Figure S2). Borneo appears to have experienced habitat restructuring rather than decline. Animations of the changes between 1700 and 2015 are provided in supplementary videos SV1and SV2.

**Figure 1.**
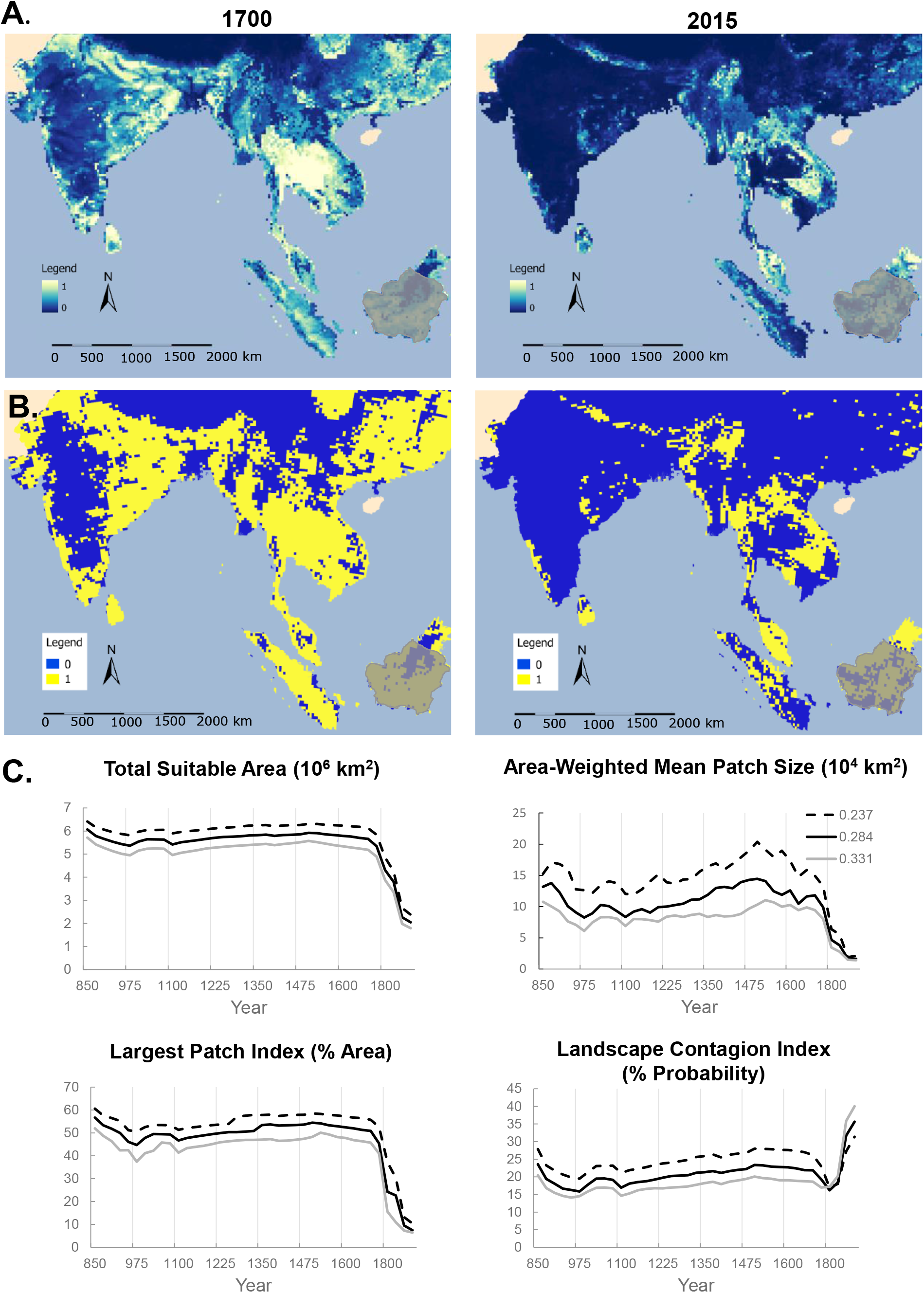
Loss of suitable habitat from 850 – 2015. Masked areas (Hainan island and part of Pakistan) have been excluded from analyses, and for visual clarity all of China is not shown. Shaded area (Borneo) is outside the currently known historic range. A) Habitat suitability predicted on the basis of elevation and the Land-use Harmonization (LUH) variables from the year 2000. B) Binarized map where 1 (yellow) indicates “suitable” areas with values above 0.284 and 0 (blue) indicates “unsuitable” areas. C) Changes in the extent and spatial configuration of suitable habitat, where each curve corresponds to the given threshold value (see methods). Total Suitable Area is the sum of all suitable habitat across the range. Area-Weighted Mean Patch Size is the weighted average of patch sizes. The Largest Patch Index is the percentage of total area occupied by the largest patch. The Landscape Contagion Index can be thought of as a measure of homogeneity, with higher probabilities representing fewer, more clumped patches. See Table S4 for complete list of fragmentation measures.

**Table 2.** Change in suitable habitat area by region from years 1700-2015. Areas are in km^2^, ordered from ranges that experienced the greatest loss to those with the greatest gain in total suitable area. “Suitable current area” refers to the current range, i.e. areas known to still contain elephants by the year 2000, whereas “Suitable potential area” refers to area that is outside the current range, where it is unknown whether elephants were ever present. The amount of suitable area within current range is 14%.

|  | Suitable Current Area (km<sup>2</sup>) |  |  | Suitable Potential Area (km<sup>2</sup>) |  |  | Total Suitable Area (km<sup>2</sup>) |  |  |
| --- | --- | --- | --- | --- | --- | --- | --- | --- | --- |
|  | 1700 | 2015 | % change | 1700 | 2015 | % change | 1700 | 2015 | % change |
| Mainland China | 1,986 | 135 | -93.2 | 1,119,258 | 65,054 | -94.2 | 1,087,183 | 65,189 | -94.2 |
| India | 216,207 | 82,793 | -61.7 | 1,439,320 | 145,561 | -89.9 | 1,655,527 | 228,354 | -86.2 |
| Bangladesh | 6,322 | 1,770 | -72.0 | 37,725 | 10,634 | -71.9 | 44,046 | 12,405 | -71.8 |
| Thailand | 40,449 | 31,303 | -22.6 | 439,964 | 127,028 | -71.2 | 480,413 | 158,331 | -67.0 |
| Vietnam | 523 | 515 | -1.4 | 196,259 | 80,923 | -58.8 | 196,781 | 81,439 | -58.6 |
| Indonesia (Sumatra) | 45,252 | 27,507 | -39.2 | 317,636 | 123,278 | -61.2 | 362,888 | 150,785 | -58.5 |
| Indonesia (Borneo) | 826 | 928 | 12.3 | 428,709 | 282,061 | -34.2 | 429,535 | 282,989 | -34.1 |
| Myanmar | 32,026 | 36,591 | 14.3 | 289,533 | 181,179 | -37.4 | 321,559 | 217,770 | -32.3 |
| Cambodia | 7,904 | 12,508 | 58.3 | 147,795 | 102,867 | -30.4 | 155,699 | 115,374 | -25.9 |
| Nepal | 12,086 | 4,750 | -60.7 | 44,333 | 37,905 | -14.5 | 56,419 | 42,655 | -24.4 |
| Sri Lanka | 31,654 | 22,603 | -28.6 | 24,097 | 19,622 | -18.6 | 55,750 | 42,225 | -24.3 |
| Bhutan | 2,033 | 1,148 | -43.6 | 5,560 | 5,126 | -7.8 | 7,593 | 6,273 | -17.4 |
| Lao PDR | 16,507 | 17,716 | 7.3 | 148,922 | 159,843 | 7.33 | 165,429 | 177,558 | 7.3 |
| Malaysia (Peninsular) | 7,796 | 10,682 | 37.0 | 95,029 | 105,418 | 10.9 | 102,825 | 116,100 | 12.9 |
| Malaysia (Borneo) | 7,216 | 12,007 | 66.4 | 100,023 | 161,316 | 61.3 | 107,239 | 173,323 | 61.6 |
| **Totals** | **428,787** | **262,956** | **-38.7** | **4,834,163** | **1,607,815** | **-66.8** | **5,228,886** | **1,870,770** | **-64.2** |

### Present-day suitability vs. distribution

Just over 48% of the current range was classified as suitable for elephants by 2015 (Figures S2 & S3, Table S3). India has the largest proportion, but only about a third was classified as suitable by the year 2015. Sri Lanka and Malaysian Borneo appear to have populations that are more than twice what would be expected relative to their share of current range, with around 63% of the range in Sri Lanka and 95% of that in Borneo qualifying as suitable (Table S3). On the other hand, Lao PDR, Thailand and Myanmar have much lower populations than expected based on their share of current range, despite approximately 79%, 60% and 51% of these ranges respectively being suitable. Most remaining range in Vietnam (98%) and Indonesian Borneo (100%) is suitable but extremely small, together accounting for just 0.27% of the total.

### Future scenarios

Suitable habitat is predicted to decline further by 2100 across all scenarios, but patterns vary by country and scenario (Figures 2, S4-S6 and supplementary video SV3). Most countries are predicted to lose habitat across the majority of scenarios, with India predicted to lose the greatest in absolute and relative terms (Figures 2 & S4). However, peninsular Malaysia may gain habitat within the current range for all cases. The Bornean portion of the range shows potential for increase in three of six scenarios. Yunnan province, China, exhibits potential habitat gains in five out of six scenarios, three of these more than doubling (Figures 2 & S4).

**Figure 2.**
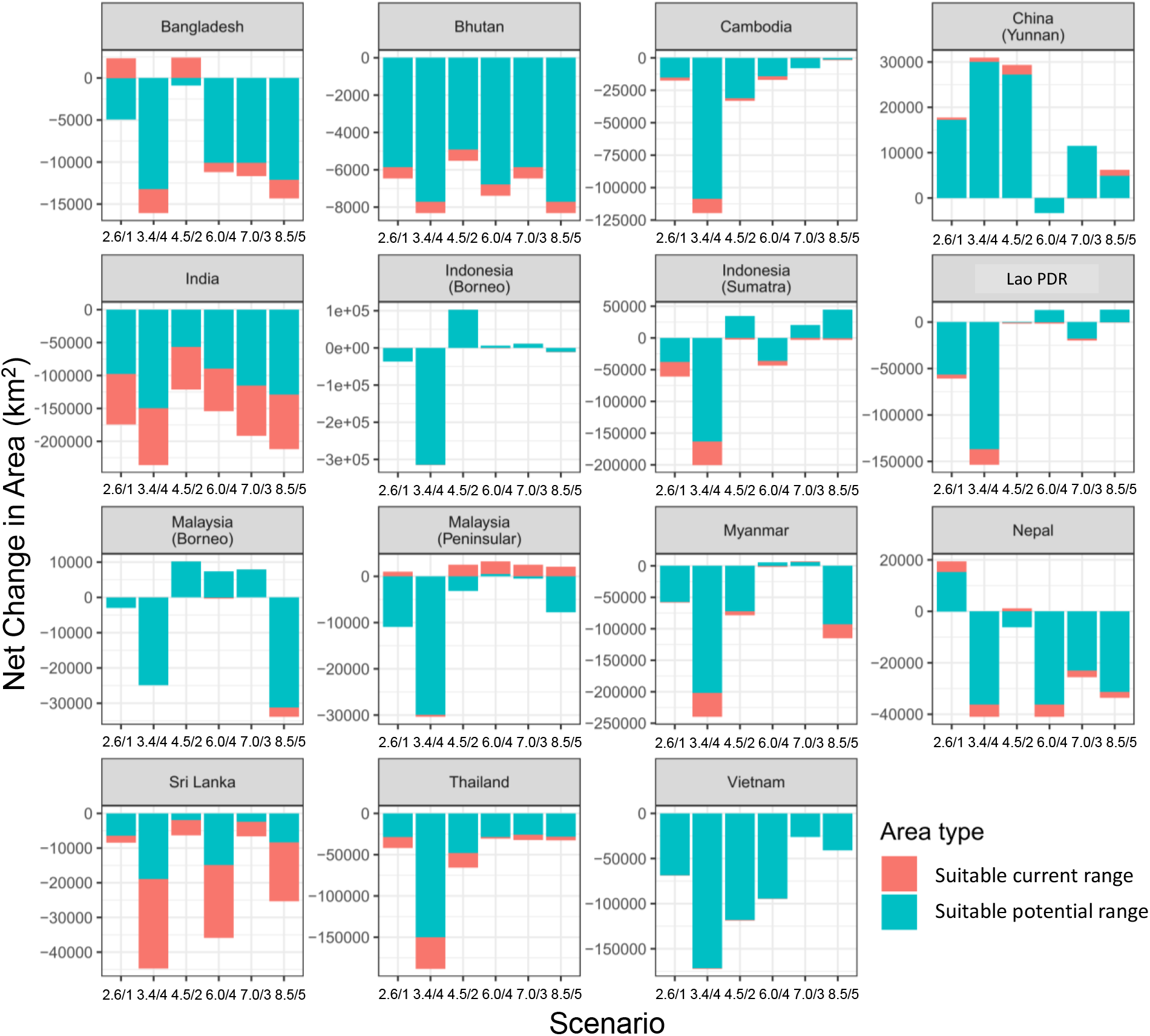
Net change in suitable habitat from 2015-2099 by country and scenario (RCP/SSP, ordered from lower to higher radiative forcings). The term “current” indicates area falling within the current range. “Suitable” indicates pixel values above the threshold of 0.284, whereas “unsuitable” indicates values below it. Increase in the “suitable current” category signifies area within the present range that was converted from unsuitable to suitable, whereas decrease signifies the reverse. These do not imply actual range expansion or contraction. “Suitable potential range” represents change in area outside the current range.

Scenarios representing lower RCPs are not consistently associated with better outcomes for these landscapes, even within countries (Figure 2 & 3, Figures S5 & S6, Table 3). Scenarios 2.6/1 and 8.5/5, with the lowest and highest amounts of radiative forcing, respectively, show similar overall losses (2.6/1 yields −24.24% and 8.5/5 yields −22.41%). Likewise, scenarios 4.5/2 and 7/3 have nearly identical changes in extent and fragmentation (Table 3, Figures 3 & S7). Strikingly, SSP4, which is associated with two different possible RCPs (3.4/4 and 6.0/4), shows greater losses in the lower RCP scenario (−83.26%) than in the higher RCP scenario (−19.29%), with 3.4/4 being the worst outcome in terms of both habitat loss and fragmentation among all six scenarios (Table 3, Figures 3 & S7). It consistently shows a severe decrease in the amount of suitable habitat accompanied by reductions in patch size and increased contagion (Figure 3 and supplementary video SV3). Other scenarios do not appreciably differ in terms of changes to metrics of habitat fragmentation and no single one appears to be better than others. Despite the overall losses, all other scenarios allow for the possibility that while decreases occur within this decade, habitats can be stabilized or partially recovered thereafter (Figures 3 & S7).

**Figure 3.**
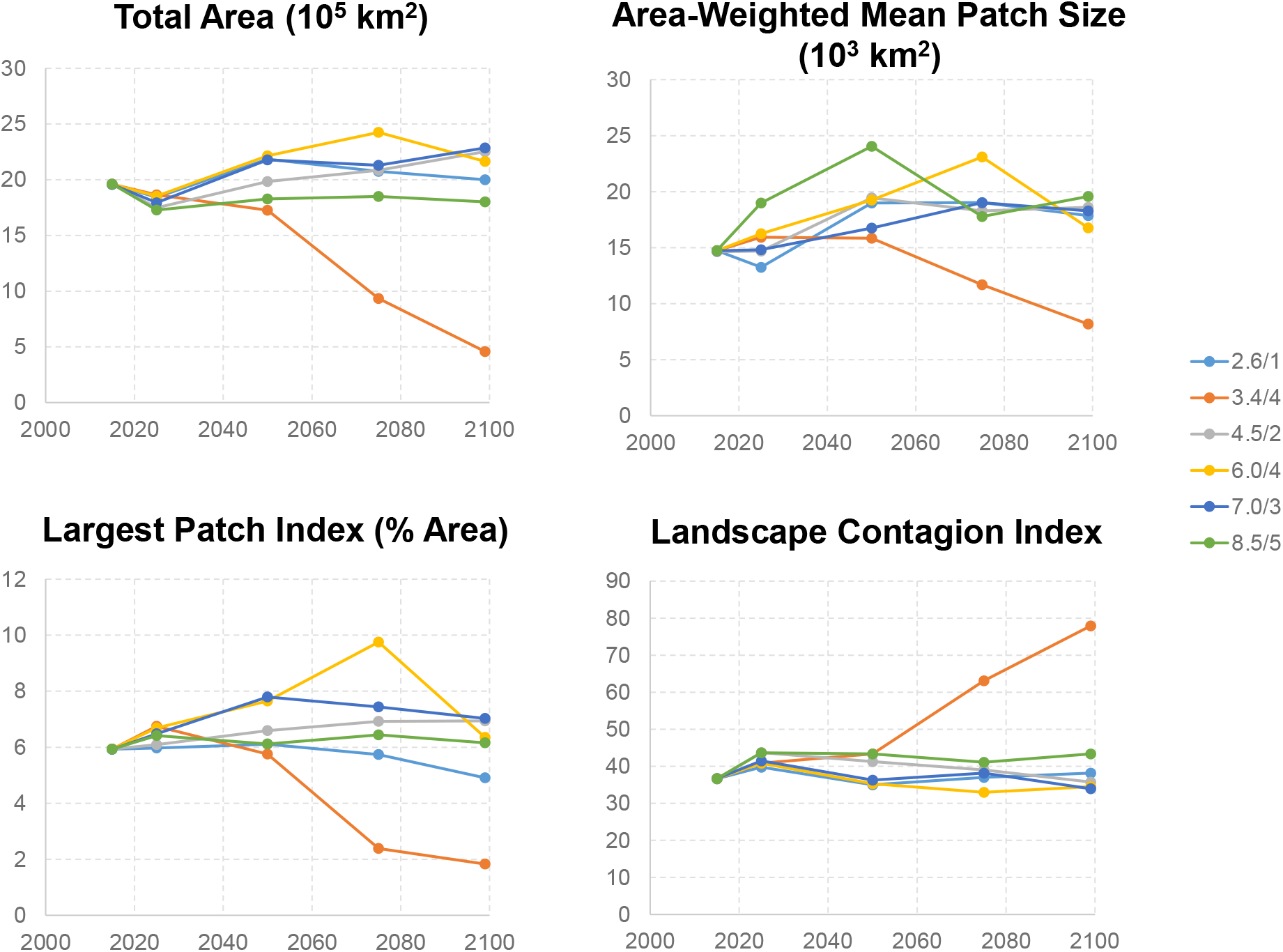
Landscape fragmentation indicators across different scenarios. No single scenario yields a consistently better outcome across all measures, however scenario 3.4/4 is consistently worse than all others.

**Table 3.** Change in suitable habitat across scenarios from 2015-2099. There is no clear association between emission levels or socioeconomic narratives and changes in habitat suitability.

| RCP/SSP | Change in suitable area within current range (%) | Change suitable area outside current range (%) | Change in total suitable area (%) |
| --- | --- | --- | --- |
| 2.6/1 | -39.2 | -21.8 | -24.2 |
| 3.4/4 | -89.5 | -82.3 | -83.3 |
| 4.5/2 | -31.1 | -9.3 | -12.3 |
| 6.0/4 | -36.0 | -16.6 | -19.3 |
| 7.0/3 | -32.0 | -9.8 | -12.8 |
| 8.5/5 | -46.0 | -18.6 | -22.4 |

## Discussion

Habitat loss due to anthropogenic activities is a global concern for numerous taxa ^2,3^. While several studies have characterized losses of particular biomes in more recent decades ^2,11^, fewer have examined long-term implications of land-use trends across diverse ecoregions. We present the first quantitative assessment of the impact of long-term land-use and global change scenarios on habitat availability for an endangered keystone species at a range-wide scale, encompassing important regional biodiversity. We discuss the processes that may drive observed results at country-level, then present overall outcomes in relation to possible climate change and socioeconomic trajectories.

We find that after several centuries of relative stability, nearly two-thirds of habitat within the 13 range countries declined within just the past 300 years. Less than half of the current range is classified as being suitable (14% of the potential habitat; Tables 2 & S3). Patch size reduction and fragmentation was far greater than is evident from recent decades alone. While it is possible that some of these trends are due to modelling error, our results closely match a review of 4,018 terrestrial mammals showing that on average 48.6% of a species’ range could be classified as “suitable” on the basis of their preferred habitats ^3^. This likely reflects a lag between land-use changes and population responses. Elephants may no longer be able to disperse into some areas, nor persist where there has been heavy offtake in the past ^23,24^. Conversely, in the absence of overharvest, they may survive for some time in suboptimal habitat even if failing to reproduce ^25^. Previous studies suggest that habitat is being lost even inside protected areas ^26^. Given that over half of the current range seems unsuitable, it may be instructive to relate levels of human-elephant conflict to our results, in addition to more fine-scaled characterizations of changes in habitat suitability. Moreover, our results reinforce that attempts to limit elephants to protected areas or extant ranges is impractical.

Trends in South Asia are largely driven by India and Sri Lanka, which contain the largest remaining wild populations (Table S3). Both countries were transformed by colonial road-building, logging, and plantations, during which elephants and other wildlife were eradicated from higher elevations and lowland rainforests ^27^. Those remaining in India now roam fragmented and largely unprotected landscapes. A study in central Assam found that HEC rates dramatically increased in the 1980s, corresponding to a drop in forest cover below 30-40% ^28^. Likewise, another study focused around the Nilgiri Biosphere reserve found that deforestation was associated with increases in negative incidents with elephants ^29^. Current range in Sri Lanka more closely matches areas classified as suitable habitat in the year 1700 than 2015 (^30^ and this paper, Figure 1). A substantial portion includes agricultural mosaics with substantial human activity ^30^, consistent with the result that anthropogenic activities and land-use types are more influential on the model than primary forests (Table S2). Yet extirpations may not occur for several generations due to the slow pace of demographic processes ^31^. Present-day development policies largely continue failing to consider the likelihood of human-elephant conflict at planning stages, followed by efforts to drive out elephants from development zones. Governmental subsidies and irrigation infrastructure promote cultivation of conflict-prone food crops such as rice, fruits and vegetables in and around remaining habitat. Both India and Sri Lanka are predicted to continue losing habitat in future scenarios (Figures 2 & S4), which will likely exacerbate conflict and drive elephant population declines, unless these trends are reversed by re-aligning development agendas with stated commitments to biodiversity conservation.

The movements of elephants and people convert ecological events into social, economic and political issues. Range shifts by elephants can introduce challenges for human communities that have little experience with elephants. For instance, while Bangladesh and Nepal can potentially gain habitat under certain scenarios, Bhutan is predicted to lose habitat under all scenarios. Yet Bhutan has a constitutional mandate to maintain at least 60% of its forest cover. All three countries share borders with India, where habitat losses are predicted and thus could experience influx of elephants. Conversely, the political displacement of people can also drastically alter habitat. In 2018 there was rapid, large-scale disruption of an elephant corridor at Cox Bazar between Bangladesh and Myanmar with the settlement of Rohingya refugees^32^. Such events might more likely under narratives SSP3 and SSP4. Future changes in political conditions as much as suitable habitat have important implications for trans-boundary management plans, wherein governments should recognize linked ecological and sociopolitical concerns.

In Southeast Asia, the disappearance of highly suitable habitat in what is now central Thailand is particularly striking (Figure 1), much of it occurring between 1950-1990 (supplementary videos SV1 and SV2). This therefore reflects not only colonial era timber extraction and land-use conversions, but also more the recent “Green Revolution” of industrial agriculture. Although expanses of forest remain in Thailand and Myanmar, both have lower estimated elephant populations than expected based on their share of current range (Table S3, Figure S4). This might be driven by habitat quality ^24^ or high rates of historic offtake for both the timber and tourism industries ^33,34^. As a result, captive elephants now outnumber wild elephants in both countries ^24,33^. Myanmar had been experiencing high rates of poaching for skins ^35^. Yet both countries are pursuing re-introduction efforts. Given the negative trends across all future scenarios, such programs might consider where suitable habitats are likely to shift to in the future (Figures S5 & S6).

Two of the most threatened elephant populations are found in Sumatra and Vietnam. Sumatra is expected to suffer losses even under low-emission scenarios. Although 98% of the current range in Vietnam is classified as suitable (Table S3), the extent of this range is extremely small and all future scenarios show losses of potential habitat outside the current range. Both cases require concerted efforts to recover habitat and re-connect isolated wildlife populations through ecosystem-level management.

Malaysia stands out as a region for potential gains both for mainland and Bornean populations, with potential connectivity^36^. In both cases, this may be at least in part because selectively logged or secondary forests have more open canopies than primary forests, encouraging the growth of understory forage. Our results support observations that secondary vegetation is more influential in predicting habitat use than primary forest (Table S2) ^37,38^. Alternatively, the model may be failing to distinguish plantation landscapes (oil palm, coffee, tea, rubber etc.) from natural forests. Because LUH2 variables explicitly differentiate primary forest from other types, the latter explanation is unlikely, but one might question how these categories were defined (see study limitations, below). Yunnan province in China may also offer opportunities for population recovery and re-wilding, if there is sufficient political desire as well as capacity to safeguard against hunting and trafficking. Most importantly, range expansion in all cases can only be possible alongside management programs designed to also safeguard human communities and livelihoods.

Surprisingly, low-emissions scenarios do not necessarily yield better outcomes than higher-emissions scenarios. This is highlighted by the similarity between scenarios 2.6/1 and 8.5/5, together with the contrast between scenarios 3.4/4 and 6.0/4 (Table 3). It is especially noteworthy that 3.4/4 resulted in distinctly worse habitat loss and fragmentation patterns than all others (Table 3, Figures 3 & S7). Policies achieving lower emissions targets strictly via advances in developed countries and/or through heavier reliance on biofuels, which already contribute heavily to habitat loss, will further endanger at-risk ecosystems. Southeast Asia contains regions with some of the greatest overlaps in carbon stocks and biodiversity but they are also among the most threatened by fuel sources such as oil palm^7,9^. Scenario 3.4/4 represents a case in which heightened regional inequities result in “a failure of current policies for energy access, leading to continued and increased use of biomass in the households of developing countries” ^39^. This underscores the complexity of the relationship between land-use practices and climate change ^6^, where climate trends cannot be divorced from societal concerns. While it is undoubtedly important to stabilize the climate by limiting global emissions, the pathway to doing so must manage Earth’s social-ecological system in a manner that does not further undermine vulnerable species, ecosystems, or human populations.^8^

### Study limitations

Habitat suitability models typically rely on relating the species of interest (i.e. occurrence, behavior) to ecological covariates. They exclude at least two important considerations. First, habitat characteristics offer a limited view of which areas may support a particular species in the absence of demographic data. Locations that animals may perceive as attractive but promote unsustainable rates of mortality are known as ecological traps.^40^ By limiting ourselves to “natural” landscapes, we have chosen to be conservative, excluding the possibility that elephants might potentially flourish in some human-dominated landscapes. This decision is underpinned by the second set of considerations: human perceptions and behavior. A species cannot flourish on a landscape that is otherwise ideal if people exclude or suppress it. Given the extent of loss in natural habitats in and around extant range, human-modified landscapes (c.f. “working landscapes”) could play a pivotal role for elephants as well as other wildlife in the future as well as re-colonized potential range, if conflicts are minimized ^19,41^.

Conversely, although we sampled occurrences from “natural” landscapes, including protected areas in which human activities are now limited, historically these biodiverse landscapes may have also been shaped by people. Traditional cultivation and diverse agroforestry practices may have been part of an unrecognized anthropological history ^42^. Communities with legacies of sustainable resource management have been displaced under unjust colonial land-use practices, including through the creation of protected areas^43^. Nevertheless, concepts such as “protected area” and “land ownership” are baked into the scenarios themselves, e.g. by assuming probability of land-use conversion to be contingent on protected status ^17,44^. These paradigms leave little room to envision other varied and socially just alternative (e.g., communal) forms of land management.

The datasets underlying our results (the LUH2 variables) are derived from a many other complex models rather than direct measurements and observations, each with assumptions and limitations. The numerous interacting components preclude directly relating results to specific features. Indeed, the scenario developers themselves refrain from commenting on ‘motivating forces,’ especially with regard to mid-range outcomes^17^. At the same time, many modelling choices entail simplification for practicality. For instance, the SSPs as a set incorporate feedbacks from climate change on biological but not economic systems ^45^. One could easily imagine such feedbacks (e.g., costly natural disasters). The longer the time horizon of a model, the greater the uncertainty and potential for error propagation. This applies equally to the niche model as much as its underlying predictors. Conditions in the past or future may represent variable combinations that do not currently occur, thus transfers between time points may involve extrapolation. For all these reasons, our outcomes are only examples of possible worlds at relatively coarse spatial resolution over long time periods rather than precise depictions of reality. Our intent is not to identify a particular “best” outcome around which policies can be crafted, nor even to predict which specific sites are of conservation value at fine-scale (see also ^46^). Rather, because Asian elephants are ecologically important flagship species for conservation management and policy with respect to diverse ecoregions, we expect these findings to prompt further discussions and studies to understand the impact of land-use trends for future planning, climate change mitigation strategies, and sustainability initiatives at national and global levels.

## Methods

### Elephant occurrence

Elephant occurrence locations were initially compiled from the Global Biodiversity Information Facility (https://www.gbif.org/), Movebank (https://www.movebank.org/) and published literature ^47–49^ as well as data contributed by the authors based on direct sightings, data logged via tracking devices, and camera traps. Records were first checked visually for errors (e.g., occurrences outside natural continental range) then refined to include locations representing natural habitat where the species could conceivably flourish. We included logged forest because secondary or regenerating forest can support elephants with potentially little conflict with humans. We excluded monoculture plantations and agricultural areas, because even if such human-dominated landscapes appear capable of supporting elephants and may even seem preferred,^41^ populations on the whole may be subject to chronic negative impacts due to accidents, conflicts with people, pollutants and gradual anthropogenic degradation of landscapes^31,50^. Moreover, we cannot know whether elephants would have in fact used human-dominated landscapes in the past when more natural areas were readily available, whereas we consider it safe to assume that elephants would have used the latter. Lastly, our goal was to quantify changes to natural biomes representing a wide array of biodiversity, without confounding these with other types of landscapes in which elephants might today persist.

To minimize sampling bias that could result in model overfitting, we subsampled data to cover the full distribution as widely as possible while eliminating redundant points located within any particular landscape. For instance, only one randomly selected data point per individual, per population, was used from any tracking dataset. Outliers from this set were removed using Cooks’ distance to eliminate locations that could represent potential errors. The final dataset consisted of 91 occurrence points spanning the years 1996-2015 which served as training data (Figure S1), where all data other than from GBIF and the literature were contributed by the authors or individuals listed in acknowledgments. QGIS and Google Earth Pro were used to visualize and consolidate these data.

### Predictor variables

The LUH2 data products were downloaded from the University of Maryland at http://luh.umd.edu/data.shtml (LUHv2h “baseline” scenario released October 14^th^ 2016 and LUHv2f future scenarios released December 12^th^ 2017). They contain three types of variables gridded at 0.25° x 0.25° (approximately 30km^2^ at the equator): state variables describing the land-use of a grid cell for a given year, transition variables describing the changes in a grid cell from one year to the next, and management variables that describe agricultural applications such as irrigation and fertilizer use, totaling 46 variables. Of these, we selected 20 variables corresponding to all three types (Table S1), which were expected to be relevant to elephant habitat use. Using ArcGIS 10 (ESRI 2017) we extracted each variable for each year between 1700-2015 from the historical reconstructions constituting LUHv2h, as well as each year between 2015-2099 for six different future scenarios from LUHv2f. We used 2099 rather than 2100 because variables representing land-use transitions, such as wood harvest rates, were not available for 2100. Note that although wood harvest rates are considered “transition” variables, in fact they represent the percentage of a pixel harvested within a given year, resulting in land-use conversions between years. We separately obtained elevation from the SRTM Digital Elevation Model (Table S5).

The RCP-SSP pairings of the LUH2 datasets (Table 1) do not include the entire space of combinations because not all outcomes are either possible or relevant. Scenarios with strong barriers to mitigation and adaptation cannot achieve low radiative forcing targets whereas those lesser barriers are only of interest with respect to the lower range of targets (see section 1 of the supplementary text) ^39^. These scenarios therefore only represent a cross-section of futures that may occur when specific socioeconomic conditions and policy decisions place differential constraints on efforts to mitigate or adapt to climate change (see supplementary text, section 1), not a factorial design.

### Data analysis

We limited the geographic extent of all analyses to the 13 range countries in which elephants are currently found. We used MAXENT, a maximum entropy algorithm^51^, to model habitat suitability. Raster files were binarized in ArcGIS into suitable and unsuitable habitat with a pixel size of approximately 20 km^2^ at a cutoff threshold (discussed below) for the years between 850 and 2000 at 25-year intervals, plus 2015. While we recognize that the suitability of any habitat patch for a species may in reality fall along a gradient, given the spatial and temporal scale of the datasets used in this study, we feel such refinements would necessitate subjective judgments that are biologically questionable. Binarization is also a practical requirement for quantifying habitat loss and fragmentation. However, as there is no commonly accepted threshold type ^52^, to ensure that the specific choice of threshold did not affect the observed trends in suitable habitat, we initially used three possible thresholds: 0.237, representing ‘maximum test sensitivity plus specificity,’ 0.284 corresponding to ‘maximum training sensitivity plus specificity,’ and 0.331 representing ‘10^th^ percentile training presence’. Unless otherwise stated, for subsequent analyses we show only results using the threshold of 0.284, where everything below this threshold was classified as ‘unsuitable’ while everything above it was classified as ‘suitable’. The binarized maps were re-projected using the WGS84 datum and an Albers Equal Area Conic projection.

To establish whether a model using the LUH2 variables yields reasonable predictions of habitat suitability for elephants, we first compared the result of using LUH2 variables for the year 2000 to one based on other features expected to directly influence present-day elephant distributions, based on knowledge of the species’ ecology ^53–55^. These include climate, terrain, land-cover, and human and livestock densities (Table S5). The details of this comparison and results are provided in the Supplementary Information (Section 3, Figures S8 & S9). The LUH2 model was then applied to all prior and future years to generate predictions of habitat suitability. Analyses were performed using the ‘dismo’ package in R (R Core Team 2017).

Polygons representing elephant range were digitized from Hedges et al. 2008 ^57^, limited to the category identified as “active confirmed”. We refer to the area within these polygons as “current range,” avoiding the term “occupied” to avoid confusion with occupancy modelling, and areas outside as “potential range”. We compared the total area of the habitat classified as suitable within and outside the current elephant range for each range country as well as Borneo and Sumatra, quantifying changes over time. We included the entirety of Borneo because both genetic studies and geological history allow for the possibility that elephants could have been natively distributed throughout the island ^58,59^, and there are no present-day physical barriers to dispersal on the island. We ranked each region based on the proportion of the elephant population found within the region as well as the percentage of current range within the region, and calculated the ratio of these ranks (Table S3). Country-level analyses were compared for all countries except China, Indonesia, and Malaysia. The extent of historic change was assessed for all of mainland China, where elephants originally occurred, but analyses of future scenarios were restricted to Yunnan province, where elephants are now confined. For Indonesia we distinguished between Sumatran and Bornean populations, in recognition of the distinct subspecies in these two regions. Likewise, for Malaysia we distinguished between Peninsular and Bornean populations.

For all regions we calculated the historic change in suitable habitat by subtracting the total area classified as suitable in the year 2015 from that of the year 1700, as major changes take place within this period (Figure 1). Likewise, we calculated the net change in projected suitable area between 2015 and 2099 across all six scenarios (Figure 2). We additionally calculated fragmentation statistics (Table S4) for all LUH2-based predictions (historical as well as future scenarios) with the program FRAGSTATS (v.4.2) ^60^. These metrics characterize changes to the spatial configuration of habitat in addition to their absolute extent. We used a ‘no sampling’ strategy and set the search radius and threshold distance, respectively, to 61 km (approximately three pixel lengths) based on the movement and dispersal capacity of elephants^61,62^.

## Acknowledgments

The authors wish to thank Kate Jenks and Megan Baker for providing some of the elephant occurrence data. AM & SdS received funding from the U.S. Fish & Wildlife Asian Elephant Conservation Fund supporting collection of elephant occurrence data in Sri Lanka and Sumatra. Preparation of this manuscript was partially supported by a James Smithson Fellowship to SdS and funding from the Section in Ecology, Behavior and Evolution at the University of California, San Diego. TW thanks the Colby College Environmental Studies Program, the Ralph J. Bunche Scholarship and Russell F. Cole Fellowship.

## Author contributions

SdS and PL conceived of the study. SdS, PL, and PN guided analyses and study design. SdS, TW, AW, AT, and JJ performed analyses. SdS and PN drafted the manuscript. All other authors contributed data to the study, revised and approved the manuscript.

## References

1. Jantz, S. M. et al. Future habitat loss and extinctions driven by land-use change in biodiversity hotspots under four scenarios of climate-change mitigation. Conserv. Biol. 29, 1122–1131 (2015).

2. Watson, J. E. M. et al. Catastrophic declines in wilderness areas undermine global environment targets. Curr. Biol. 26, 2929–2934 (2016).

3. Crooks, K. R. et al. Quantification of habitat fragmentation reveals extinction risk in terrestrial mammals. Proc. Natl. Acad. Sci. 114, 7635–7640 (2017).

4. IPBES. Global assessment report on biodiversity and ecosystem services of the Intergovernmental Science-Policy Platform on Biodiversity and Ecosystem Services. (IPBES Secretariat, 2019).

5. Lambin, E. F., Geist, H. J. & Lepers, E. Dynamics of land-use and land-cover change in tropical regions. Annu. Rev. Environ. Resour. 28, 205–241 (2003).

6. Díaz, S. et al. Pervasive human-driven decline of life on Earth points to the need for transformative change. Science. 366, (2019).

7. Soto-Navarro, C. et al. Mapping co-benefits for carbon storage and biodiversity to inform conservation policy and action. Philos. Trans. R. Soc. B Biol. Sci. 375, (2020).

8. O’Neill, D. W., Fanning, A. L., Lamb, W. F. & Steinberger, J. K. A good life for all within planetary boundaries. Nat. Sustain. 1, 88–95 (2018).

9. Ripple, W. J. et al. Collapse of the world’s largest herbivores. Sci. Adv. 1, e1400103 (2015).

10. Tucker, M. A. et al. Moving in the Anthropocene: Global reductions in terrestrial mammalian movements. Science. 359, 466–469 (2018).

11. Hansen, M. C. et al. High-resolution global maps of 21 st-century forest cover change. Science (80-.). 342, 850–853 (2013).

12. Beller, E. E., McClenachan, L., Zavaleta, E. S. & Larsen, L. G. Past forward: Recommendations from historical ecology for ecosystem management. Glob. Ecol. Conserv. 21, (2020).

13. Strassburg, B. B. N., Iribarrem, A. & Beyer, H. L. Global priority areas for ecosystem restoration. Nature (2020). doi:10.1038/s41586-020-2784-9

14. Rondinini, C. et al. Global habitat suitability models of terrestrial mammals. Philos. Trans. R. Soc. B Biol. Sci. 366, 2633–2641 (2011).

15. Soberón, J. & Townsend Peterson, A. Monitoring biodiversity loss with primary species-occurrence data: toward national-level indicators for the 2010 target of the Convention on Biological Diversity. AMBIO A J. Hum. Environ. 38, 29–34 (2009).

16. Sánchez-Cordero, V., Illoldi-Rangel, P., Linaje, M., Sarkar, S. & Peterson, A. T. Deforestation and extant distributions of Mexican endemic mammals. Biol. Conserv. 126, 465–473 (2005).

17. O’Neill, B. C. et al. The roads ahead: Narratives for shared socioeconomic pathways describing world futures in the 21st century. Glob. Environ. Chang. 42, 169–180 (2015).

18. Campos-Arceiz, A. & Blake, S. Megagardeners of the forest – the role of elephants in seed dispersal. Acta Oecologica 37, 542–553 (2011).

19. Nyhus, P. J. Human-Wildlife Conflict and Coexistence. Annual Review of Environment and Resources 41, (2016).

20. Olivier, R. Distribution and status of the Asian Elephant. Oryx 14, 379–424 (1978).

21. Hurtt, G. C. et al. Harmonization of Global Land Use Change and Management for the Period 850-2100 (LUH2) for CMIP6. Earth System Grid Federation (2020).

22. Moss, R. H. et al. The next generation of scenarios for climate change research and assessment. Nature 463, 747–756 (2010).

23. Calabrese, A. et al. Conservation status of Asian elephants: the influence of habitat and governance. Biodivers. Conserv. 26, 2067–2081 (2017).

24. Songer, M., Aung, M., Allendorf, T. D., Calabrese, J. M. & Leimgruber, P. Drivers of change in Myanmar’s wild elephant distribution. Trop. Conserv. Sci. 9, (2016).

25. de Silva, S. & Leimgruber, P. Demographic tipping points as early indicators of vulnerability for slow-breeding megafaunal populations. Front. Ecol. Evol. 7, 1–13 (2019).

26. Neupane, D., Kwon, Y., Risch, T. S. & Johnson, R. L. Changes in habitat suitability over a two decade period before and after Asian elephant recolonization. Glob. Ecol. Conserv. 22, e01023 (2020).

27. Skinner, T. Fifty years in Ceylon: an autobiography. (Tisara Press, 1891).

28. Chartier, L., Zimmermann, A. & Ladle, R. J. Habitat loss and human-elephant conflict in Assam, India: does a critical threshold exist? Oryx 45, 528–533 (2011).

29. Puyravaud, J.-P., Gubbi, S., Poornesha, H. C. & Davidar, P. Deforestation increases frequency of incidents with elephants (Elephas maximus). Trop. Conserv. Sci. 12, 194008291986595 (2019).

30. Fernando, P., de Silva, M. K. C. R., Jayasinghe, L. K. A., Janaka, H. K. & Pastorini, J. First country-wide survey of the Endangered Asian elephant: towards better conservation and management in Sri Lanka. Oryx 1–10 (2019). doi:10.1017/s0030605318001254

31. de Silva, S. & Leimgruber, P. Demographic tipping points as early indicators of vulnerability for slow-breeding megafaunal populations. Front. Ecol. Evol. 7, (2019).

32. Goldsmith, B. Elephants face ‘time bomb’ in Bangladesh land clash with Rohingya refugees. (2019).

33. Jackson, J., Childs, D. Z., Mar, K. U., Htut, W. & Lummaa, V. Long-term trends in wild-capture and population dynamics point to an uncertain future for captive elephants. Proceedings. Biol. Sci. 286, 20182810 (2019).

34. Nijman, V. An assessment of the live elephant trade in Thailand. (2014).

35. Sampson, C. et al. New elephant crisis in Asia — Early warning signs from Myanmar. PLoS One 13, e0194113 (2018).

36. de la Torre, J. A. et al. Using elephant movements to assess landscape connectivity under Peninsular Malaysia’s central forest spine land use policy. Conserv. Sci. Pract. 1, e133 (2019).

37. Evans, L. J., Goossens, B., Davies, A. B., Reynolds, G. & Asner, G. P. Natural and anthropogenic drivers of Bornean elephant movement strategies. Glob. Ecol. Conserv. 22, e00906 (2020).

38. Rood, E., Ganie, A. A. & Nijman, V. Using presence-only modelling to predict Asian elephant habitat use in a tropical forest landscape: Implications for conservation. Divers. Distrib. 16, 975–984 (2010).

39. Riahi, K. et al. The Shared Socioeconomic Pathways and their energy, land use, and greenhouse gas emissions implications: An overview. Glob. Environ. Chang. 42, 153–168 (2017).

40. Robertson, B. & Hutto, R. A framework for understanding ecological traps and an evaluation of existing evidence. Ecology 87, 1075–1085 (2006).

41. de la Torre, J. A. et al. There will be conflict – agricultural landscapes are prime, rather than marginal, habitats for Asian elephants. Anim. Conserv. 24, 720–732 (2021).

42. Cairns, M. Shifting Cultivation Policies: Balancing Environmental and Social Sustainability. (2017).

43. Rangarajan, M. & Shahabuddin, G. Displacement and relocation from protected areas: Towards a biological and historical synthesis. Conserv. Soc. 4, 359–378 (2006).

44. Hasegawa, T., Fujimori, S., Ito, A., Takahashi, K. & Masui, T. Global land-use allocation model linked to an integrated assessment model. Sci. Total Environ. 580, 787–796 (2017).

45. Calvin, K. et al. The SSP4: A world of deepening inequality. Glob. Environ. Chang. 42, 284–296 (2017).

46. Di Marco, M., Watson, J. E. M., Possingham, H. P. & Venter, O. Limitations and trade-offs in the use of species distribution maps for protected area planning. J. Appl. Ecol. 54, 402–411 (2017).

47. Hedges, S. et al. Distribution, status, and conservation needs of Asian elephants (Elephas maximus) in Lampung Province, Sumatra, Indonesia. Biol. Conserv. 124, 35–48 (2005).

48. Santiapillai, C. & Dissanayake, S. R. B. Observations on elephants in the Maduru Oya National Park Sri Lanka (Mammalia, Elephantidae). Gajah 20, 9–20 (2001).

49. Marasinghe, M. S. L. R. P., Dayawansa, N. D. K. & Silva, R. P. Area suitability prediction for conserving elephants: An application of likelihood ratio prediction model. Trop. Agric. Res. 25, (2015).

50. Goswami, V. R., Vasudev, D. & Oli, M. K. The importance of conflict-induced mortality for conservation planning in areas of human–elephant co-occurrence. Biol. Conserv. 176, 191–198 (2014).

51. Hijmans, R. J. & Elith, J. Species distribution modeling with R. (2013).

52. Bean, W. T., Stafford, R. & Brashares, J. S. The effects of small sample size and sample bias on threshold selection and accuracy assessment of species distribution models. Ecography (Cop.). 35, 250–258 (2012).

53. Fernando, P. & Leimgruber, P. Asian elephants and seasonally dry forests. in The Ecology and Conservation of Seasonally Dry Forests in Asia (eds. McShea, W. J., Davies, S. J., Phumpakphan, N. & Pattanavibool, A.) 151–163 (Smithsonian Insitution Scholarly Press, 2011).

54. Matawa, F., Murwira, A. & Schmidt, K. S. Explaining elephant (Loxodonta africana) and buffalo (Syncerus caffer) spatial distribution in the Zambezi Valley using maximum entropy modelling. Ecol. Modell. 242, 189–197 (2012).

55. Pastorini, J. et al. A preliminary study on the impact of changing shifting cultivation practices on dry season forage for Asian elephants in Sri Lanka. Trop. Conserv. Sci. 6, 770–780 (2013).

56. Team, R. D. C. R: A language and environment for statistical computing. (2017).

57. Hedges, S., Fisher, K. & Rose, R. Range-wide mapping workshop for Asian elephants (Elephas maximus). (2008).

58. Sharma, R. et al. Genetic analyses favour an ancient and natural origin of elephants on Borneo. Sci. Reports2 8, 1–11 (2018).

59. Husson, L., Boucher, F. C., Sarr, A. C., Sepulchre, P. & Cahyarini, S. Y. Evidence of Sundaland’s subsidence requires revisiting its biogeography. J. Biogeogr. 47, 843–853 (2020).

60. McGarigal, K., Cushman, S. A. & Ene, E. FRAGSTATS v4: Spatial pattern analysis program for categorical and continuous maps. (2012).

61. Moßbrucker, A. M. Modeling the fate of Sumatran elephants in Bukit Tigapuluh Indonesia: research needs & implications for population management. J. For. Sci. 10, 5–18 (2016).

62. Fernando, P. et al. Ranging behavior of the Asian elephant in Sri Lanka. Mamm. Biol. - Zeitschrift für Säugetierkd. 73, 2–13 (2008).

