## Supplementary material for "The Past, Present and Future of Elephant Landscapes in Asia"

### Supplementary Text, Tables and Figures

### 1. Understanding Representative Concentration Pathways (RCPs) and Shared Socioeconomic Pathways (SSPs)

#### *1.1 A very brief introduction to RCPs*

The representative concentration pathways are a set of modelled scenarios based on emission, concentration and land-use trajectories, originally released in 2011. They are of global extent, gridded at 0.5 x 0.5 degree resolution. Our description here is based on an overview given by van Vuuren et al. 2011:

> "The word '*representative*' signifies that each of the RCPs represents a larger set of scenarios in the literature. In fact, as a set, the RCPs should be compatible with the full range of emissions scenarios available in the current scientific literature, with and without climate policy. The words '*concentration pathway*' are meant to emphasize that these RCPs are not the final new, fully integrated scenarios (i.e. they are not a complete package of socio-economic, emission and climate projections), but instead are internally consistent sets of projections of the components of radiative forcing that are used in subsequent phases. The use of the word '*concentration*' instead of '*emissions*' also emphasizes that concentrations are used as the primary product of the RCPs, designed as input to climate models."

The first thing of note above is that these scenarios are not exhaustive, but rather a subset taken to be *representative* of the vast literature. They rest on models developed by various independent research groups, distilling work conducted by a very large community (van Vuuren, Edmonds, et al., 2011).

Some terms to disambiguate are *concentration*, *emissions*, and *radiative forcing*. The term "concentration" refers to the concentration of all greenhouse gases in the atmosphere, not only emissions. Radiative forcing (or climate forcing) is a measure of the rate of change in energy per unit area, measured in watts per square meter ($W/m^2$) that is the result of both natural (e.g. fires) and anthropogenic processes (e.g. emissions, deforestation) thus they are also not solely a measure of emissions, although there is clearly a positive relationship between the two. The various RCP labels refer to radiative forcing levels achieved by the end of the timeframe under consideration, the year 2100, with discrete scenario outcomes ranging from 2.6-8.5 $W/m^2$.

The key drivers underlying all models consist of UN population projections, the 90$^{th}$ percentile range of GDP scenarios in the literature on greenhouse gas emissions, energy use by type of sector (i.e. fossil fuels, biofuels, renewables), land-use scenarios, emissions scenarios, and the concentrations of greenhouse gases and air pollutants. However, each have their own distinct assumptions and policy interventions, therefore they are not directly comparable to one another nor should outcomes from different scenarios be meaningfully averaged together. Moreover, note that there can be multiple ways of achieving a particular RCP and therefore the achieved target is not necessarily a unique outcome of a particular modelling framework.

#### *1.2 A very brief introduction to SSPs*

Although the RCPs derive from internally-consistent socio-economic assumptions, as a set they "do not provide an internal logic nor do they span the full range of socio-economic trajectories in the literature" (van Vuuren et al. 2011). The Shared Socioeconomic Pathways framework provides this structure (O'Neill et al., 2014, 2015; Riahi et al., 2017). The SSPs are described as "plausible alternative trends in the evolution of society and natural systems over the 21st century at the level of the world and large world regions (O'Neill et al., 2014)." These are a set of five narratives (colorfully described as types of 'roads') accompanied by quantitative measures of development. The socioeconomic drivers are population growth, economic activity and urbanization, in turn yielding a wide range of outcomes in terms of energy use, emissions and landuse. The SSPs characterize scenarios that fall along two axes pertaining to socioeconomic challenges for mitigation and socioeconomic challenges for adaptation (O'Neill et al., 2015):

| | | Challenges to adaptation | | |
|---|---|---|---|---|
| | | **Low** | **Moderate** | **High** |
| Challenges to mitigation | **Low** | SSP 1: *Sustainability – taking the green road* | | SSP 5: *Fossil-fueled development – Taking the highway* |
| | **Moderate** | | SSP 2: *Middle of the road* | |
| | **High** | SSP 4: *Inequality – A road divided* | | SSP 3 *Regional Rivalry – A rocky road* |

No scenario offers a particularly close representation of the present or future, as each one contains elements that are observable in current society as well as imaginings about the future (O'Neill et al., 2015). Importantly, these scenarios represent conditions that constrain the degree to which a society can achieve a given radiative forcing target; model runs with high mitigation or adaptation challenges are unable to achieve low radiative forcing targets. Thus the possible futures that could obtain are themselves envisioned as a matrix, being the combinations of RCPs along one axis and SSPs along the other, but not all RCP-SSP pairings are possible (O'Neill et al., 2014).

#### *1.3 The RCP-SSP Pairings of the LUH2 Datasets*

The following are brief characterizations of the scenarios of the LUH2 datasets with some details relevant to the present study. For fuller descriptions, please refer to Hurtt et al. 2020 and references therein.

**RCP 2.6-SSP1**

This pairing is based on IMAGE, the Integrated Model to Assess the Global Environment 3.0 and considers the potential for keeping global mean temperature increase below 2$^{o}$C through the combined effects of agricultural and energy systems, land cover, carbon and hydrological cycles and climate change (Stehfest et al., 2014). SSP1 is characterized as a "green" development paradigm, where population growth levels off mid-century with a high degree of environmentally-conscious economic growth and technological improvements. Negative emissions from energy use in the second half of the 21st century are to be achieved through a combination of bio-energy, reforestation and carbon capture and storage as well as improvements in energy efficiency. Of note is the statement that this decrease is "slightly offset by an increase in land-use related $CO_2$ emissions compared to baseline due to use of land for bioenergy production" (van Vuuren, Stehfest, et al., 2011). Relevant for our purposes, according to van Vuuren et al. 2011, land use is allocated on the basis of rules pertaining to the potential for agricultural productivity, proximity to existing agricultural areas, proximity to current water bodies, and a random factor. From this it is possible that areas in close proximity to existing areas of industrial-scale biofuel production (e.g. parts of southeast Asia) may be more likely to continue and even expand such activities. However, an important feature of SSP1 is cooperation and technology transfer among regions, with more developed countries providing access to human, financial and technological resources to those that are developing (O'Neill et al., 2015).

**RCP 3.4-SSP4**

This pairing derives from GCAM, the Global Change Assessment Model (Wise et al., 2014) and represents an attempt to implement strong climate change mitigation policies imposing a high carbon tax within a context of extremely heterogeneous global development (Calvin et al., 2017; O'Neill et al., 2015). The schism in global society is driven by a combination of uneven investments in human capital, together with the consolidation of political power among social and economic elites, widening the gap between the "haves" and "have nots". High and middle-income countries experience economic growth and increase their energy demands, which are met through diversification including renewable and nuclear technologies as well as non-traditional biofuels. However lower income countries struggle to meet basic health and sanitation needs, with consumption driven by increased population growth. These regions have weaker institutions and governments have a lesser degree of control over land-use. Inefficiencies in agricultural production foster the huge increase in cultivated land. Consequently, there is afforestation in middle- and high-income countries but deforestation in low-income countries.

**RCP 4.5-SSP2**

This pairing derives from the IIASA Integrated Assessment Modelling (IAM) framework. Within it, land-use is modelled through the Global Biosphere Management Model or GLOBIOM with spatially explicit projections made through G4M and then linked to the MESSAGE energy system model (Hurtt et al., 2020). Hurtt et al. 2020 state nothing about the actual land-use features of this particular implementation of RCP4.5, aside from the following: "An important feature of RCP4.5 is the initial decrease in forest by about 43 million ha from 2000 to 2050 (comparable to the reference scenario), with a subsequent increase in forest by about 331 million ha from 2050 to 2100." SSP2 is provided as one possible middle-of-the-road pathway without socioeconomic extremes, but also without any suggestion that is more or less likely than scenarios that do envision such extremes (O'Neill et al., 2015). Development follows a trajectory that is not dissimilar to historic trends, with progress being uneven among countries, but economies and political systems are relatively stable on the whole. Demographic transitions are completed by the latter half of the century, with population growth levelling off.

**RCP 6-SSP4**

This pairing derives from the same conditions as RCP 3.4-SSP4, except that climate policies are less stringent.

**RCP 7-SSP3**

This pairing derives from a combination of AIM and Computable General Equilibrium (AIM/GCE) model that has been integrated into a land-use allocation model (Hasegawa et al., 2017). The land-use allocation is implemented through profit-maximization decisions by land owners at a local scale. The amount of pasture land is scaled according to demand, and assigned to areas that were unprotected and unused for either crop production or afforestation. It has elements of SSP4, such as high levels of global income inequality and weak institutions, but these manifest politically as tendencies toward authoritarianism and more local autonomy as opposed to the control of resources by global elites. It also differs in being less oriented toward biofuels or traditional forms of bioenergy and more reliant on fossil-fuels, similar to SSP5.

**RCP 8.5-SSP5**

This pairing is derived from the Regionalized Model of Investment and Development (REMIND) and the Model of Agricultural Production and its Impacts on the Environment (MAgPIE) baseline of SSP5. It has the highest level of energy consumption out of all five scenarios, primarily fossil-fuels, concurrent with a doubling of global food demand and tripling of greenhouse gas emissions (Hurtt et al., 2020). There is also a heavy increase in the need for concentrated livestock feed. These result in "strong expansion of global cropland into pasture and forest land, with an increase of about 300 Mha (20 %) between 2010 and 2100" (Hurtt et al., 2020).

### 2. Supplementary Figures and Tables

**Figure S1 - Presumed Asian elephant range contraction.** Brown shaded region shows presumed historic post-glacial range (Olivier 1978), smaller purple polygons show current range (classified as "active confirmed" in Hedges et al. 2008), points show sampled occurrences.

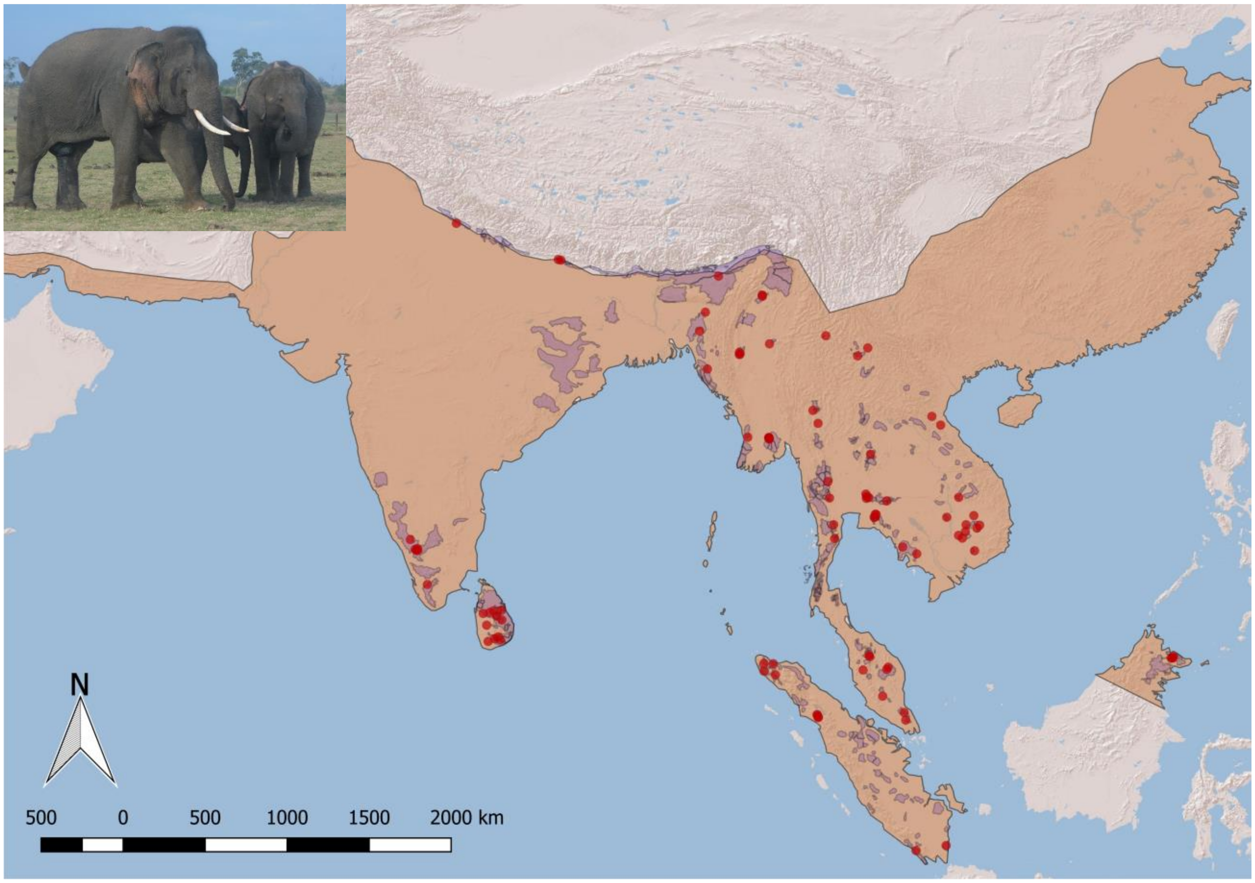

**Figure S2 – Loss and gain in suitable habitat across the range between 1700-2015.** Masked areas have been excluded from analysis. Overall, 64.2 % of the total area converted from suitable to unsuitable in this period, with 38.6 % occurring within the current range (Table 3). Habitat gains largely occurred outside the current range.

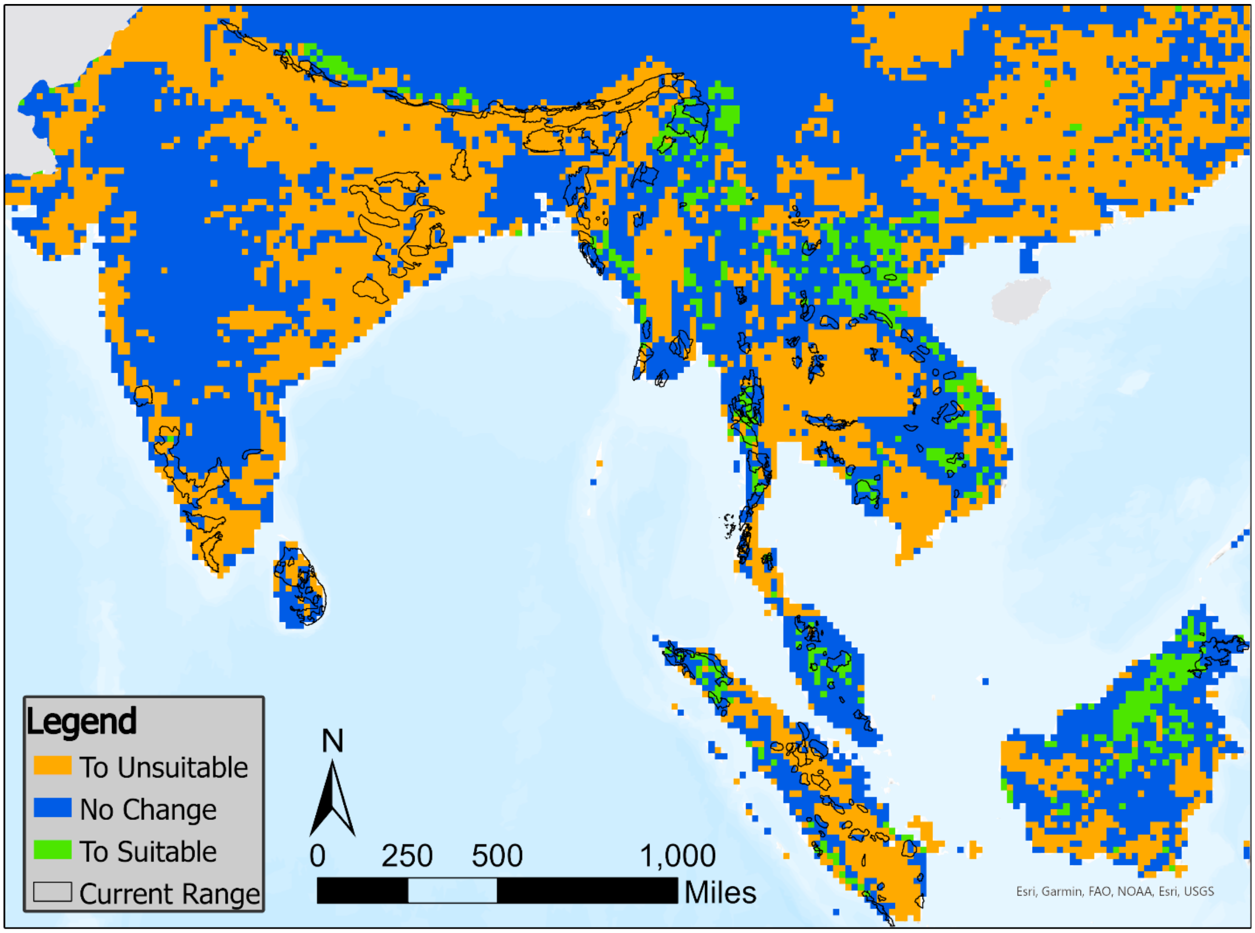

**Figure S3** – Suitability of range in the year 2015 for areas representing the current range and buffer areas of 25, 50 or 100 km. Yellow pixels indicate suitable areas, blue pixels indicate unsuitable areas. Upper panel was binarized by the threshold of maximum training sensitivity plus specificity (0.284), lower panel was binarized by the threshold of $10^{th}$ percentile training presence (0.331).

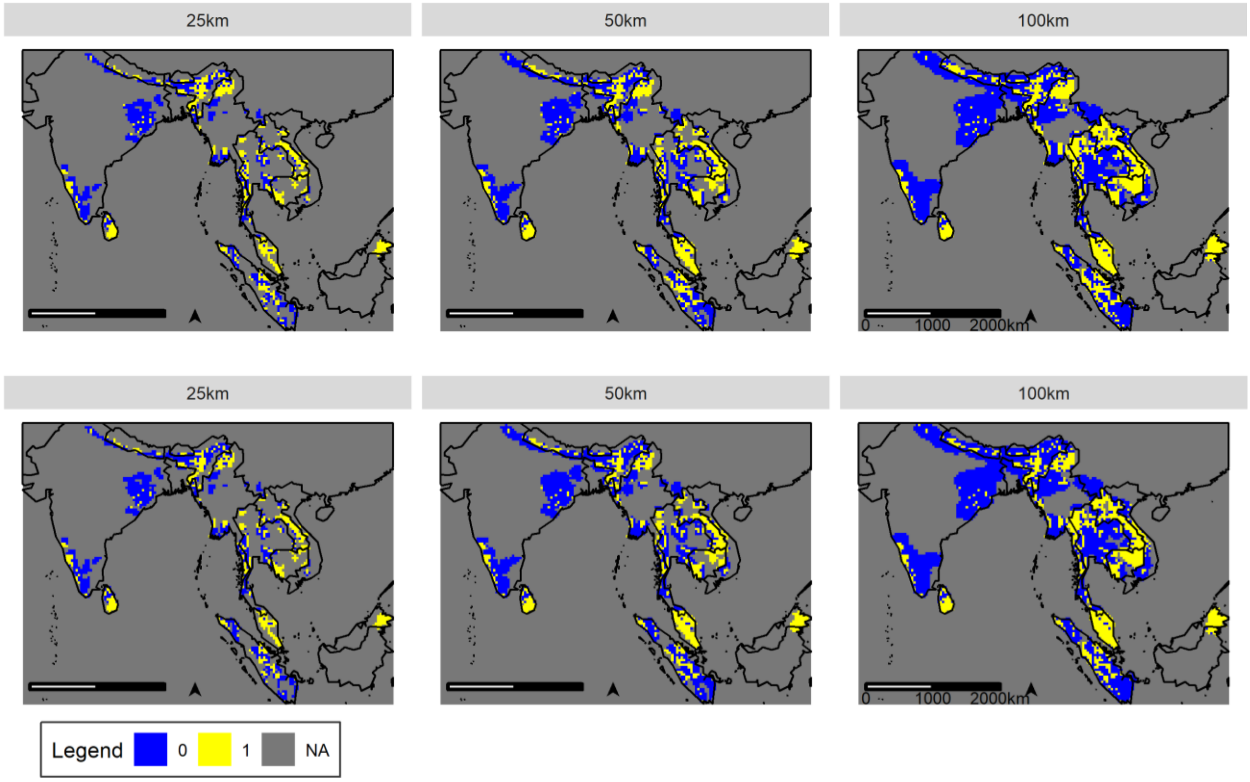

**Figure S4 – Change in suitable habitat relative to current range and population size.** The x-axis shows the fraction of land area within each region that constitutes present-day elephant range, whereas the y-axis shows the ratio of change between 2015-2100. Bubbles are scaled relative to the elephant population size. Colors represent the size of the elephant population relative to available area, with “High” being a rank ratio >1.75, “Balanced” being a ratio of 0.76-1.75, and “Low” being a ratio of 0-0.75 (Table S3). 1 = Bangladesh; 2 = Bhutan; 3 = Cambodia; 4 = China (Yunnan); 5 = India; 6 = Indonesia (Borneo); 7 = Indonesia (Sumatra); 8 = Lao PDR; 9 = Malaysia (Borneo); 10 = Malaysia (Peninsular); 11 = Myanmar; 12 = Nepal; 13 = Sri Lanka; 14 = Thailand; 15 = Vietnam.

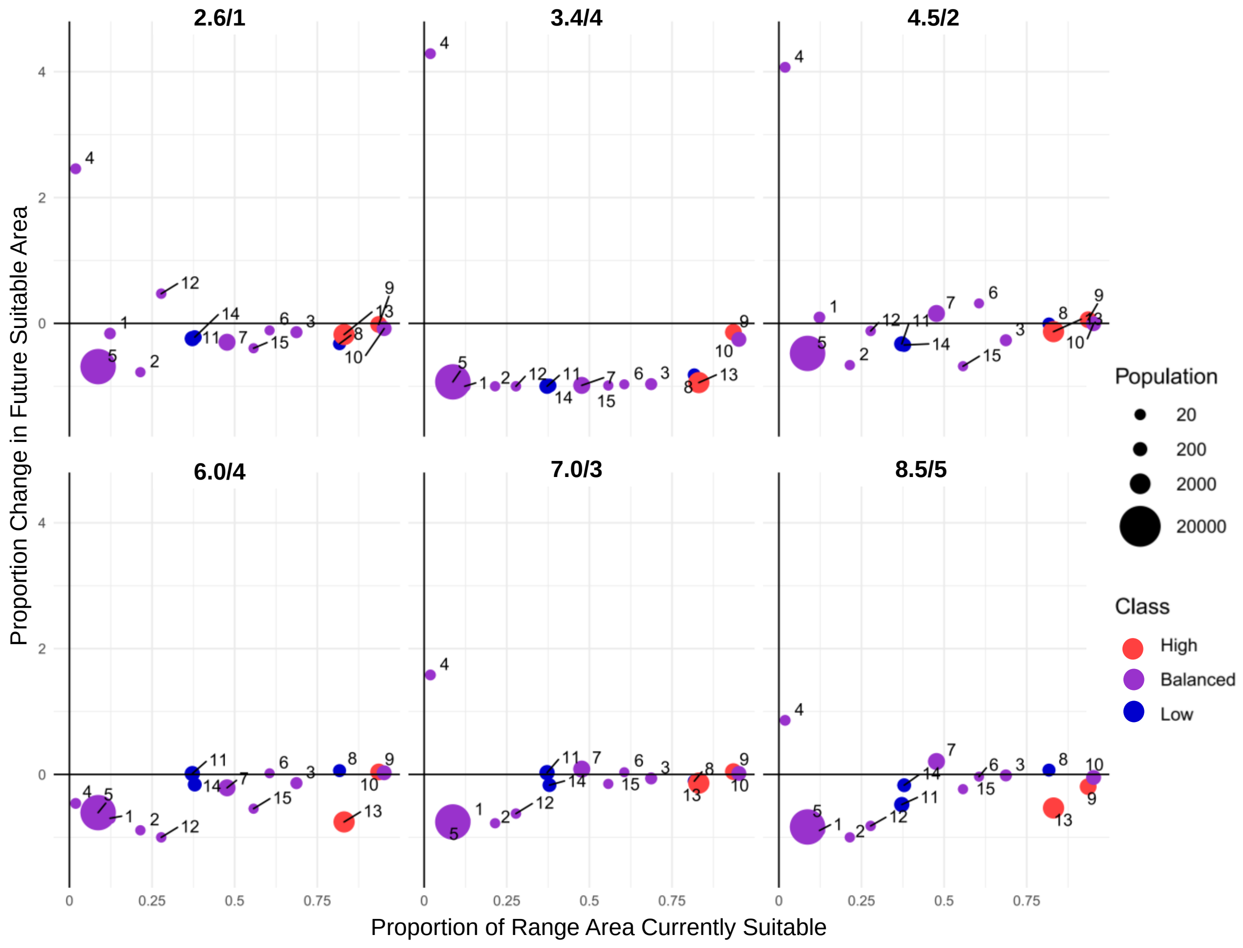

**Figure S5 – Suitability by 2099 predicted under the six different scenarios at a threshold of 0.284.** Upper panel shows area including current range and 25 km buffer, lower panel shows a 100km buffer.

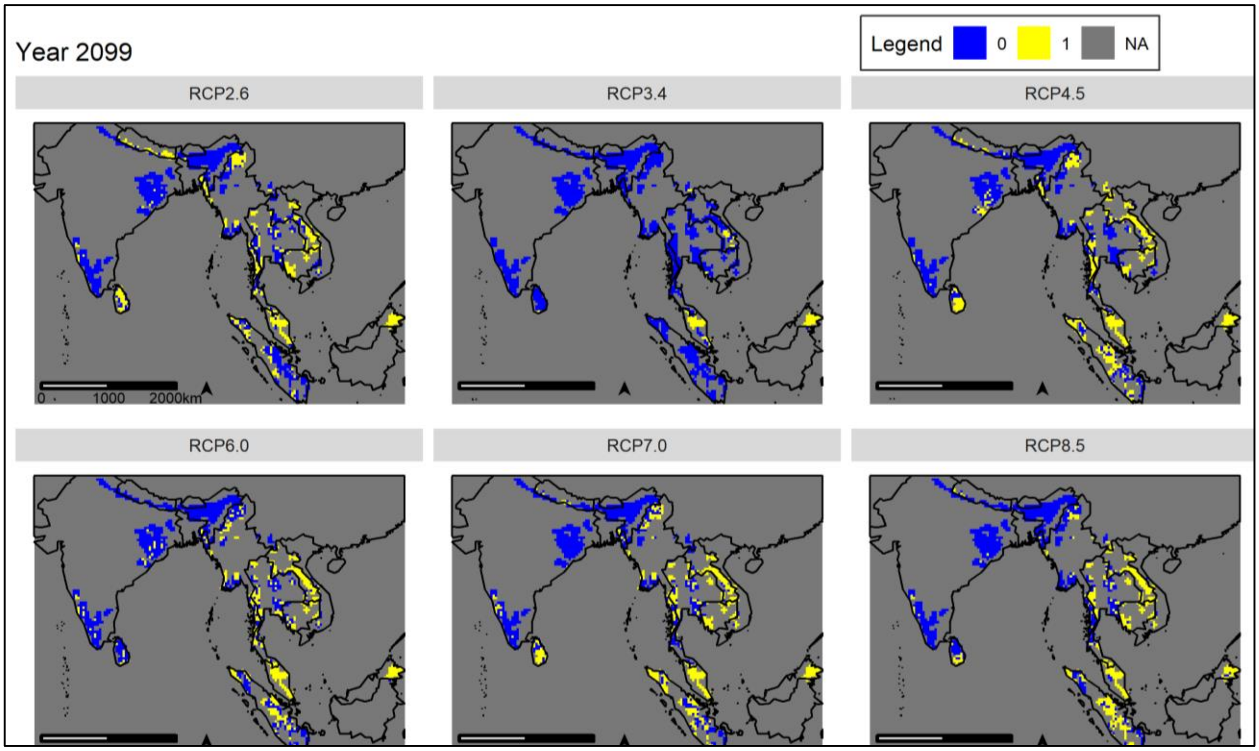


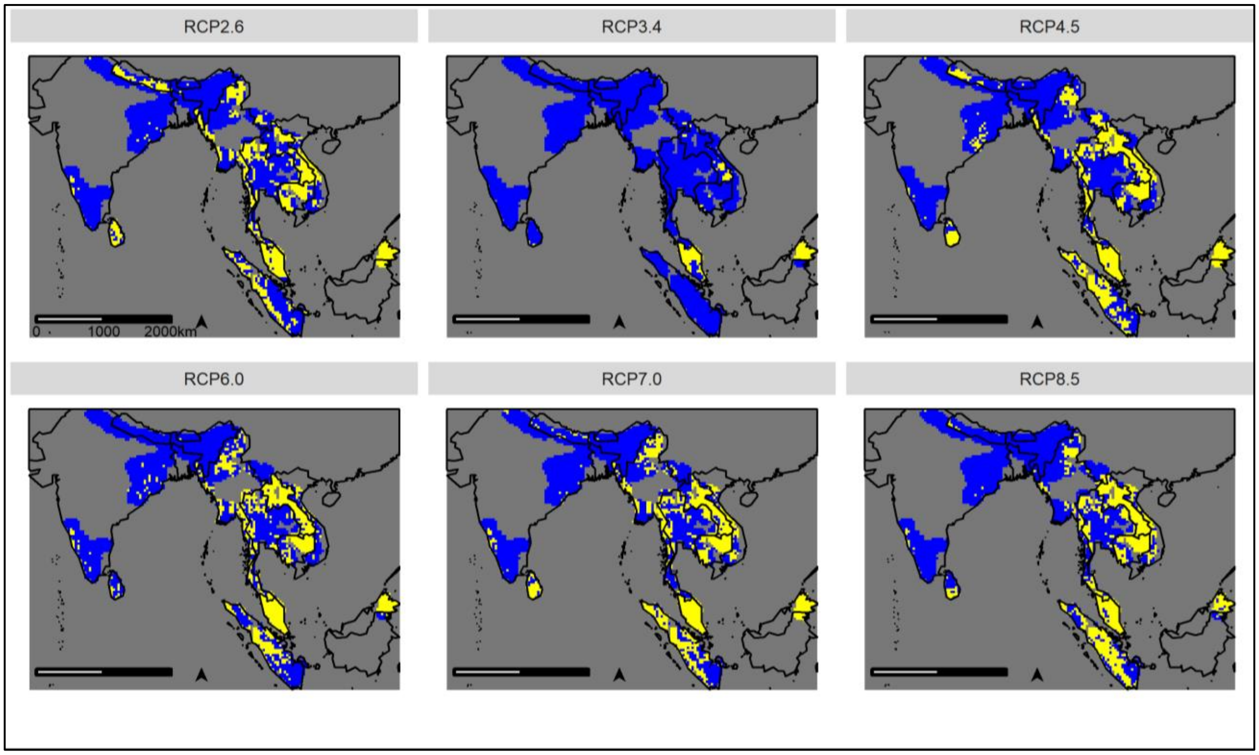

**Figure S6 – Suitability by 2099 predicted under the six different scenarios at a threshold of 0.331.** Upper panel shows area including current range and 25 km buffer, lower panel shows a 100km buffer.

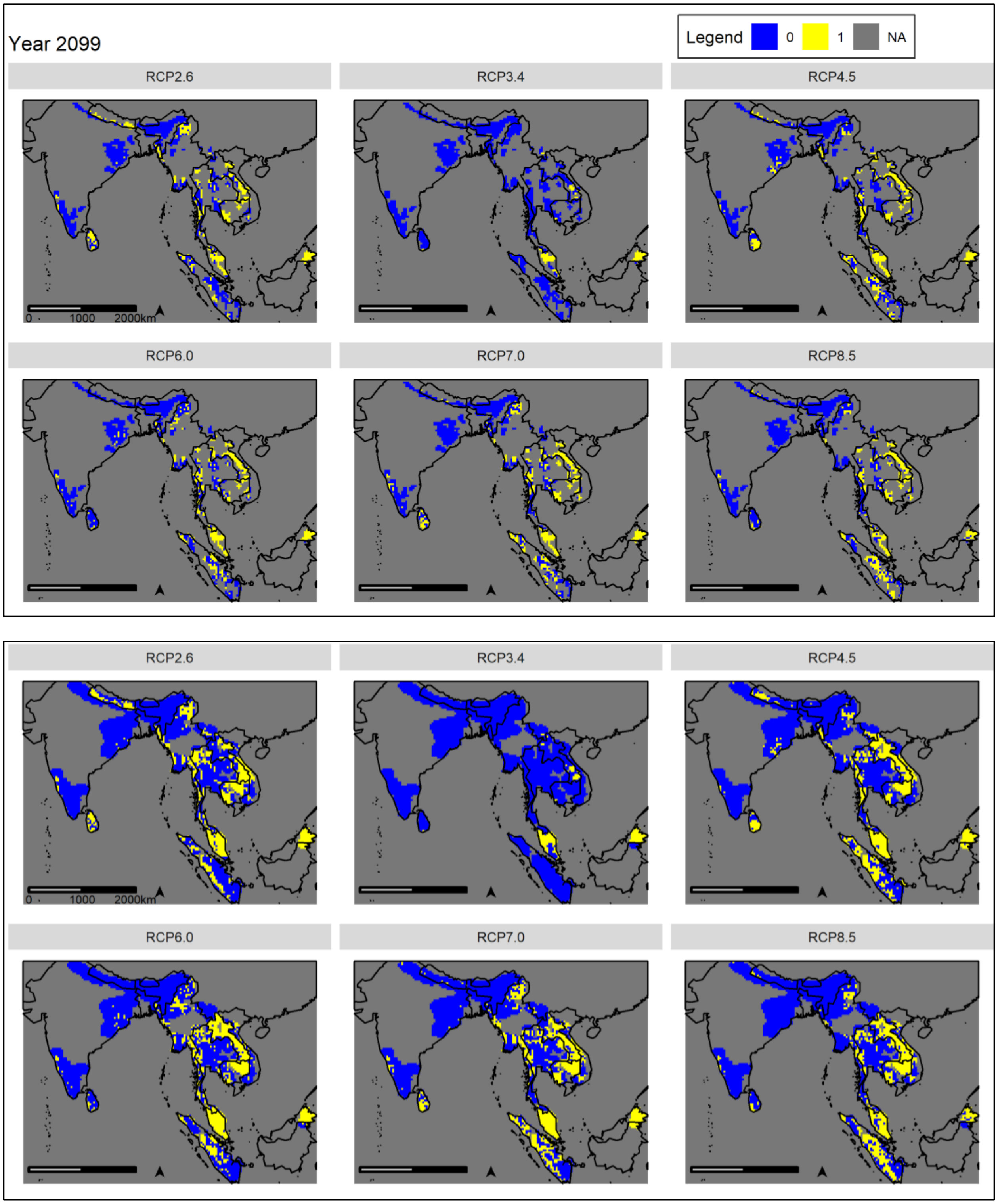

**Figure S7 – Fragmentation patterns under future scenarios under the 10th percentile training presence threshold of 0.331.** The higher threshold reduces all patch sizes resulting in a rescaling of axes compared to the results in the main text. Most notably, the area-weighted mean patch size drops by an order of magnitude. Nevertheless, the overall trends remain virtually indistinguishable from the lower threshold (main text, Figure 3).

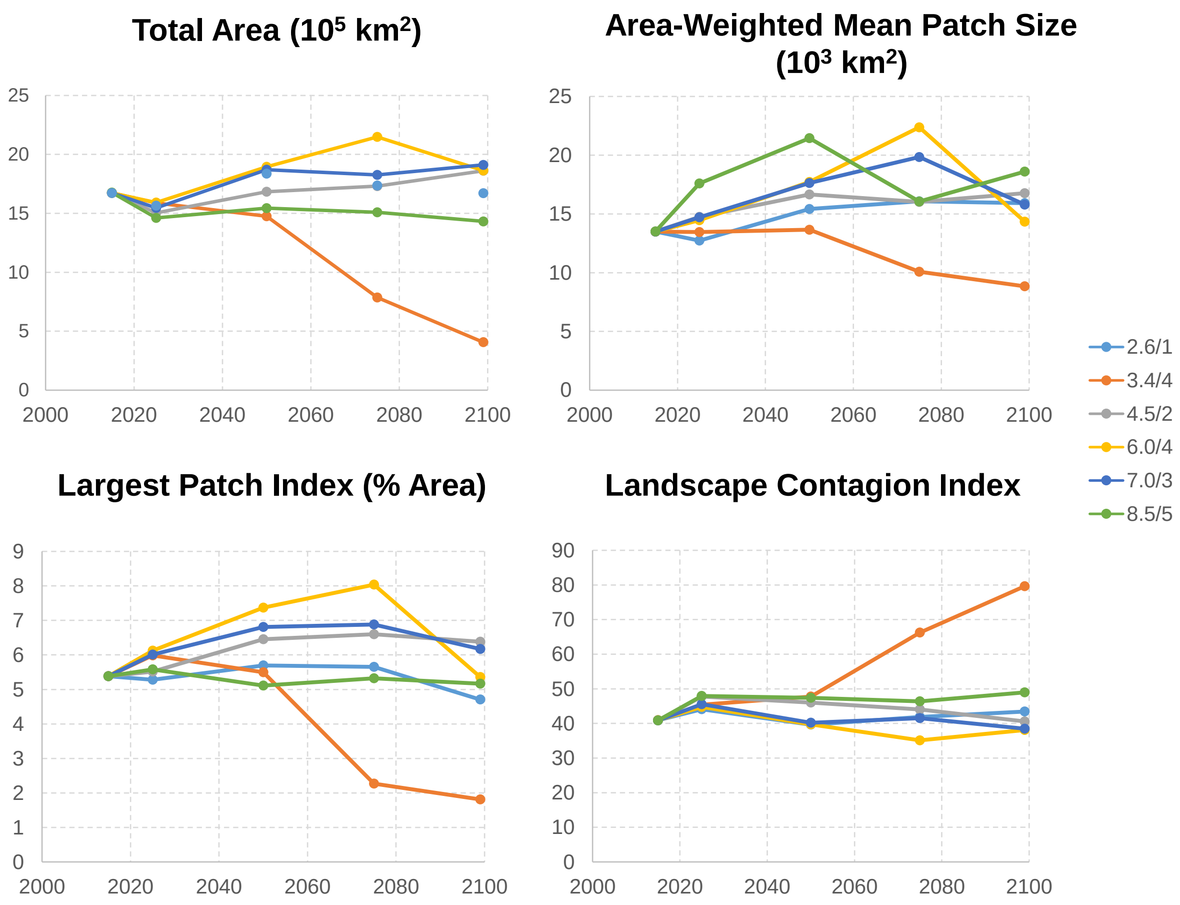

**Table S1 – Selected Land-Use Harmonization (LUH) variables.**

| States (units: fraction of grid cell) | Transitions between land use states (units: fraction of grid cell per year) | Management – Irrigation (units: fraction of crop area) |
|---|---|---|
| primf: forested primary land | primf_harv: wood harvest area from primary forest | irrig_c3ann: irrigated fraction of C3 annual area |
| primn: non-forested primary land | primn_harv: wood harvest area from primary non-forest | irrig_c3per: irrigated fraction of C3 perennial area |
| secdf: potentially forested secondary land | secmf_harv: wood harvest area from secondary mature forest | irrig_c4ann: irrigated fraction of C4 annual area |
| secdn: potentially non-forested secondary land | secyf_harv: wood harvest area from secondary young forest | irrig_c4per: irrigated fraction of C4 perennial area |
| pastr: managed pasture | | irrig_c3nfx: irrigated fraction of C3 nitrogen-fixing area |
| c3ann: C3 annual crops | | flood: flooded fraction of C3 annual crop area |
| c3per: C3 perennial crops | | |
| c4ann: C4 annual crops | | |
| c4per: C4 perennial crops | | |
| c3nfx: C3 nitrogen-fixing crops | | |

**Table S2 – LUH2 predictors with relative contributions >1%.** Variables are ordered from most to least influential, noting that their contribution can be driven by either positive or negative associations. See supplementary Table S1 for complete lists of variables.

| Variable | Variable contribution (% change in AUC) | Permutation importance |
|---|---|---|
| C3 nitrogen-fixing crops | 29.8 | 5.6 |
| SRTM digital elevation | 17.3 | 12.8 |
| Potentially non-forested secondary land | 9.9 | 10.7 |
| Non-forested primary land | 8.2 | 9.8 |
| C3 annual crops | 8.1 | 16.3 |
| C4 perennial crops | 6.5 | 4.9 |
| Managed pasture | 5.1 | 8.5 |
| C3 perennial crops | 4.1 | 0.3 |
| Wood harvest area from secondary mature forest | 3.2 | 5.8 |
| C4 annual crops | 2.1 | 1.8 |
| Primary forest | 1.9 | 5.9 |

**Table S3 – Elephant population sizes relative to available range.** "Total current range" refers to the extent of range in which elephants were thought to be present in the 2000s. "Current suitable 2015" refers to the amount of this current range still classified as suitable habitat by the year 2015 under the LUH2 model. Ranges are ranked by the percentage of the global elephant population found within them as well as the percentage of global range they encompass. They are ordered by rank ratio, which is the area rank divided by the population rank. A ratio close to 1 indicates that the population size is proportional to the amount of area within that range, higher ratios indicate populations are larger than expected on the basis of available range, and lower ratios indicate the opposite (See also Figure S4).

| Range | Wild Elephant Population | % Pop. In Range | Total Current Range ($km^2$) | % Of Current Range | Current Suitable ($km^2$) | % Current Suitable 2015 | Range Rank, Population | Range Rank, Area | Rank Ratio |
|---|---|---|---|---|---|---|---|---|---|
| Sri Lanka | 5,879 | 13.4 | 36,196 | 6.7 | 22,603 | 62.5 | 2 | 5 | 2.50 |
| Malaysia (Borneo) | 2,268 | 5.1 | 12,589 | 2.3 | 12,007 | 95.4 | 4 | 9 | 2.25 |
| China (Yunnan) | 186 | 0.4 | 2,362 | 0.4 | 135 | 5.7 | 11 | 13 | 1.18 |
| Malaysia (Peninsular) | 1,450 | 3.3 | 13,413 | 2.5 | 10,682 | 79.6 | 6 | 7 | 1.17 |
| Indonesia (Borneo) | 167 | 0.4 | 928 | 0.2 | 928 | 100 | 12 | 14 | 1.17 |
| Bangladesh | 325 | 0.7 | 6,770 | 1.3 | 1,770 | 26.1 | 10 | 11 | 1.10 |
| India | 27,000 | 61.4 | 239,056 | 44.1 | 82,793 | 34.6 | 1 | 1 | 1.00 |
| Indonesia (Sumatra) | 2,600 | 5.9 | 56,033 | 10.4 | 27,507 | 49.1 | 3 | 3 | 1.00 |
| Vietnam | 97 | 0.2 | 527 | 0.1 | 515 | 97.7 | 15 | 15 | 1.00 |
| Cambodia | 425 | 1.0 | 12,975 | 2.4 | 12,508 | 96.4 | 9 | 8 | 0.89 |
| Bhutan | 105 | 0.2 | 2,424 | 0.5 | 1,148 | 47.4 | 14 | 12 | 0.86 |
| Nepal | 126 | 0.3 | 12,178 | 2.3 | 4,750 | 39.0 | 13 | 10 | 0.77 |
| Lao PDR | 700 | 1.6 | 22,494 | 4.2 | 17,716 | 78.8 | 8 | 6 | 0.75 |
| Thailand | 1,000 | 2.3 | 52,415 | 9.7 | 31,303 | 59.7 | 7 | 4 | 0.57 |
| Myanmar | 1,619 | 3.7 | 71,281 | 13.1 | 36,591 | 51.3 | 5 | 2 | 0.40 |
| **Totals** | 43,947 | 100 | 541,640 | 100 | 262,956 | 48.6 | - | - | - |

[a]From Fernando & Pastorini, 2011. Note that these population estimates reflect the time frame relevant to the datasets used in analyses rather than the most current estimates, which may have changed.
[b]Calculated from Hedges et al. 2008 (Figure S1).

**Table S4. Summary of variables analyzed in FRAGSTATS.** Only results for total area, area-weighted mean patch size, largest patch index, and contagion are shown as these most clearly and intuitively illustrate trends in habitat suitability and structure (Figure 1 main text).

| Patch metrics | Class metrics | | Land metrics | |
|---|---|---|---|---|
| Patch area (AREA) | Total Area (CA/TA) | | Contagion (CONTAG) | |
| Euclidean Nearest-Neighbor Distance (ENN) | Percentage of Landscape (PLAND) | | IJI | |
| | Largest Patch Index (LPI) | | Proximity Index (PROX_?) | MN |
| | | | | AM |
| | | | | CV |
| | Patch Area (AREA_?) | Mean (MN) | CONNECT | |
| | | Area-Weighted Mean (AM) | | |
| | | Median (MD) | | |
| | | Range (RA) | | |
| | | Coefficient of Variation (CV) | | |
| | Perimeter-Area Fractal Dimension (PARFRAC) | | | |
| | Contiguity Index (CONTIG_?) | MN | | |
| | | AM | | |
| | | Standard Deviation (SD) | | |
| | | CV | | |
| | Euclidean Nearest Neighbor Distance (ENN_?) | MN | | |
| | | AM | | |
| | | CV | | |
| | Number of Patches (NP) | | | |
| | Interspersion Juxtaposition Index (IJI) | | | |
| | Proximity Index (PROX_?) | MN | | |
| | | AM | | |
| | | CV | | |
| | Connectance Index (CONNECT) | | | |

### 3. Comparison of habitat suitability predictions using Land-Use Harmonization (LUH) variables vs. other contemporary benchmark variables.

#### *2.1 Data analyses*

As a sanity check, we first compared the suitability map derived from LUH variables for the year 2000 (Table S1) to one derived from other features, which we refer to as *benchmark variables* (Table S2). We used the year 2000 LUH variables because it is near the midpoint of the timeframe over which elephant occurrence data were available. The benchmark variables were resampled to match the variable with the coarsest resolution (0.5°) before constructing the MAXENT model. Raster files were then binarized in ArcMap into suitable and unsuitable habitat for both sets of variables, with a cutoff threshold corresponding to 'maximum training sensitivity plus specificity' (see methods). For the benchmark model this value was 0.350, for the LUH model it was 0.284; everything below the threshold was thus classified as 'unsuitable' while everything above was classified as 'suitable' for subsequent analyses. In order to compare the results derived from both sets of predictors, the results of the benchmark variables were again resampled to match the coarser resolution of the LUH dataset.

**Table S5. Benchmark environmental predictor variables.** Contribution and permutation importance for the MAXENT model are listed for those variables with relative contributions >1% change in AUC, with variables ordered from most to least influential.

| Variable | Variable contribution (% change in AUC) | Permutation importance | Time period used | Spatial Resolution | Source |
|---|---|---|---|---|---|
| Bioclim seasonality | 46.9 | 41.4 | 2000 | ~1 km | http://www.worldclim.org/current (Hijmans, 2005) |
| FAO sheep and goat density | 16 | 15.4 | 2005 | 0.05 Decimal Degree, ~5 km | http://www.fao.org/geonetwork/srv/en/main.search?extended=off&remote=off&any=glw+12717&themekey=&to=&from=&siteId=&hitsPerPage=10 (Robinson, 2014) |
| Percent tree cover | 15 | 0.2 | 2000 | 250 m | Matt Hansen, University of Maryland https://earthengine.google.org/#detail/UMD%2Fhansen%2Fglobal_forest_change_2013 |

| | | | | | |
|---|---|---|---|---|---|
| Earthstat cropland | 5.4 | 7.1 | 2000 | 5 min (~10 km) | http://www.earthstat.org/data-download/ (Ramankutty, 2008) |
| SRTM digital elevation | 4.6 | 7.1 | 2003 | 1 km | http://www.cgiar-csi.org/data/srtm-90m-digital-elevation-database-v4-1#citation (Jarvis, 2008) |
| Landscan human population | 4.2 | 2.8 | 2009 | 1 km | http://web.ornl.gov/sci/landscan/landscan_documentation.shtml (Vijayaraj, 2008) |
| Earthstat pasture | 3.5 | 0.9 | 2000 | 5 min (~10km) | http://www.earthstat.org/data-download/ (Ramankutty, 2008) |
| Annual mean temperature | 2 | 2.3 | 2000 | ~1 km | http://www.worldclim.org/current (Hijmans, 2005) |
| Percent non-vegetated cover | 1.9 | 21.8 | 2001 | 250 m | https://lpdaac.usgs.gov/dataset_discovery/modis/modis_products_table/mod44b (DiMiceli, 2011) |
| Slope | | | 2003 | 1 km | Derived from SRTM DEM |
| Percent Non-Vegetated Cover | | | 2001 | 250 m | https://lpdaac.usgs.gov/dataset_discovery/modis/modis_products_table/mod44b (DiMiceli 2011) |
| Percent Non-Tree Vegetation | | | 2001 | 250 m | https://lpdaac.usgs.gov/dataset_discovery/modis/modis_products_table/mod44b (DiMiceli, 2011) |
| FAO Cattle Density | | | 2005 | 0.05 Decimal Degree, ~5 km | http://www.fao.org/geonetwork/srv/en/main.search?extended=off&remote=off&any=glw+12713&themekey=&to=&from=&siteId=&hitsPerPage=10 (Robinson, 2014) |

#### *3.2 Results of comparison*

We first compared predictions of habitat suitability under the benchmark model to those under the LUH model (Figure S1). After binarization, datasets were in agreement for 80% of pixels overall (>7 x $10^6$ $km^2$) and 89% of pixels within the current elephant range. They were in agreement on over 80% of the area for 6 out of 13 countries (China, Bangladesh, India, Sri Lanka, Bhutan and Cambodia; Figure S2). Indonesia, Myanmar, Thailand and Vietnam had lower levels of agreement. The lowest agreement occurred for Indonesia (58% for the Sumatran area, 56% for the Bornean area) and Vietnam (53% of area). Areas of disagreement tended to occur at intermediate values, i.e., at the transition zones from suitable to unsuitable, reflecting loss of information in converting from graded to binary outputs. Slightly more pixels were classified as “suitable” under the benchmark model relative to the LUH model (Figure S1D), thus the latter was more conservative. Nevertheless, given the high levels of agreement, we considered the LUH model reliable.

**Figure S8 – Comparison of habitat suitability modelled with the benchmark vs. LUH variables.** In (A) and (B) blue represents less suitable areas whereas yellow represents more suitable areas. The masked areas (island of Hainan and part of Pakistan) were not included in analyses. C) 1=Pixel classified as unsuitable under both models; 2=Pixel classified as suitable under benchmark model and unsuitable under LUH model; 3=Pixel classified as unsuitable under benchmark model and suitable under LUH; 4=Pixel classified as suitable under both models. D) Histogram of pixel classifications in (C).

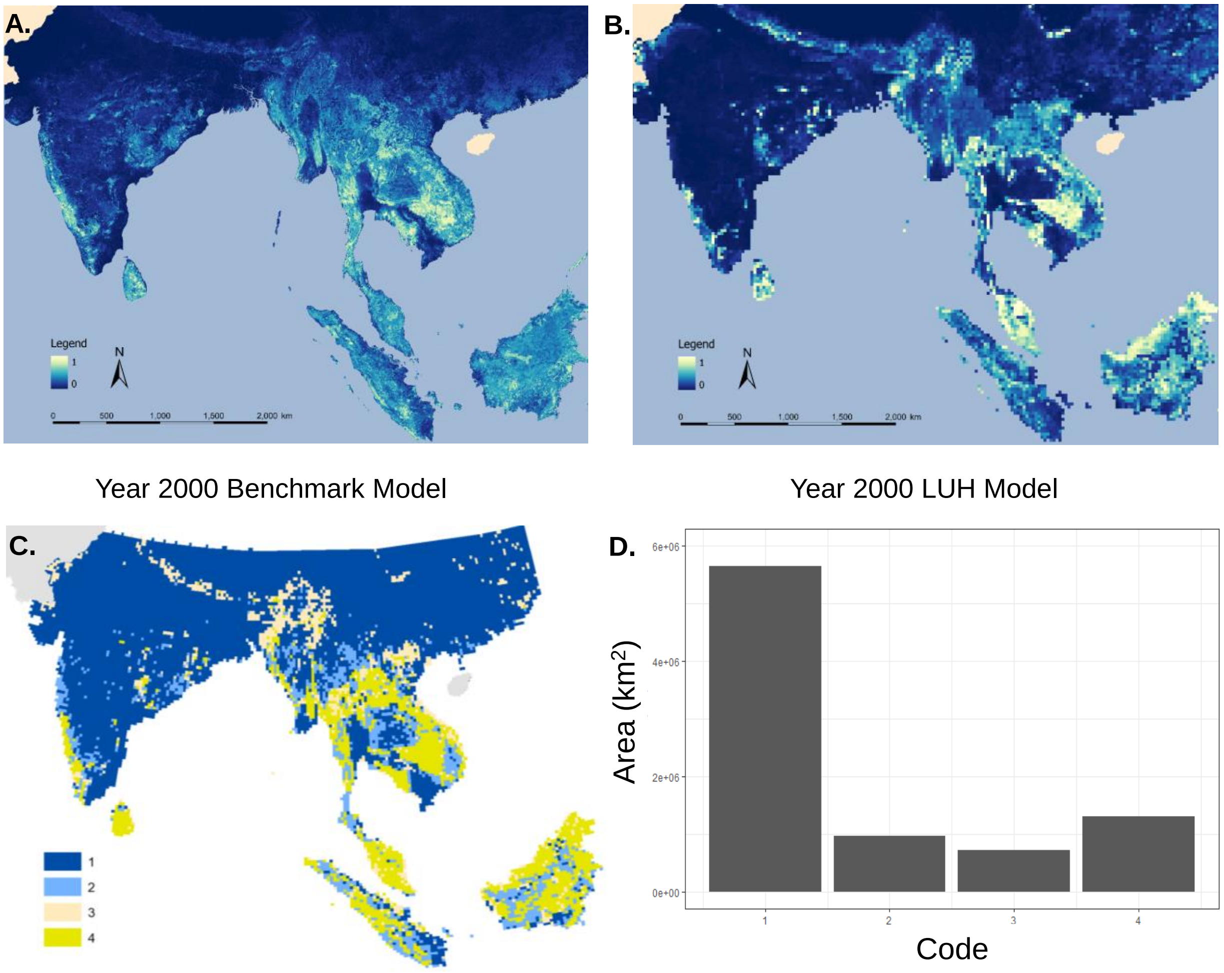

**Figure S9 – Degree of agreement between LUH and benchmark models upon binarization.** 1=Pixel classified as unsuitable under both models; 2=Pixel classified as suitable under benchmark model and unsuitable under LUH model; 3=Pixel classified as unsuitable under benchmark model and suitable under LUH; 4=Pixel classified as suitable under both models.

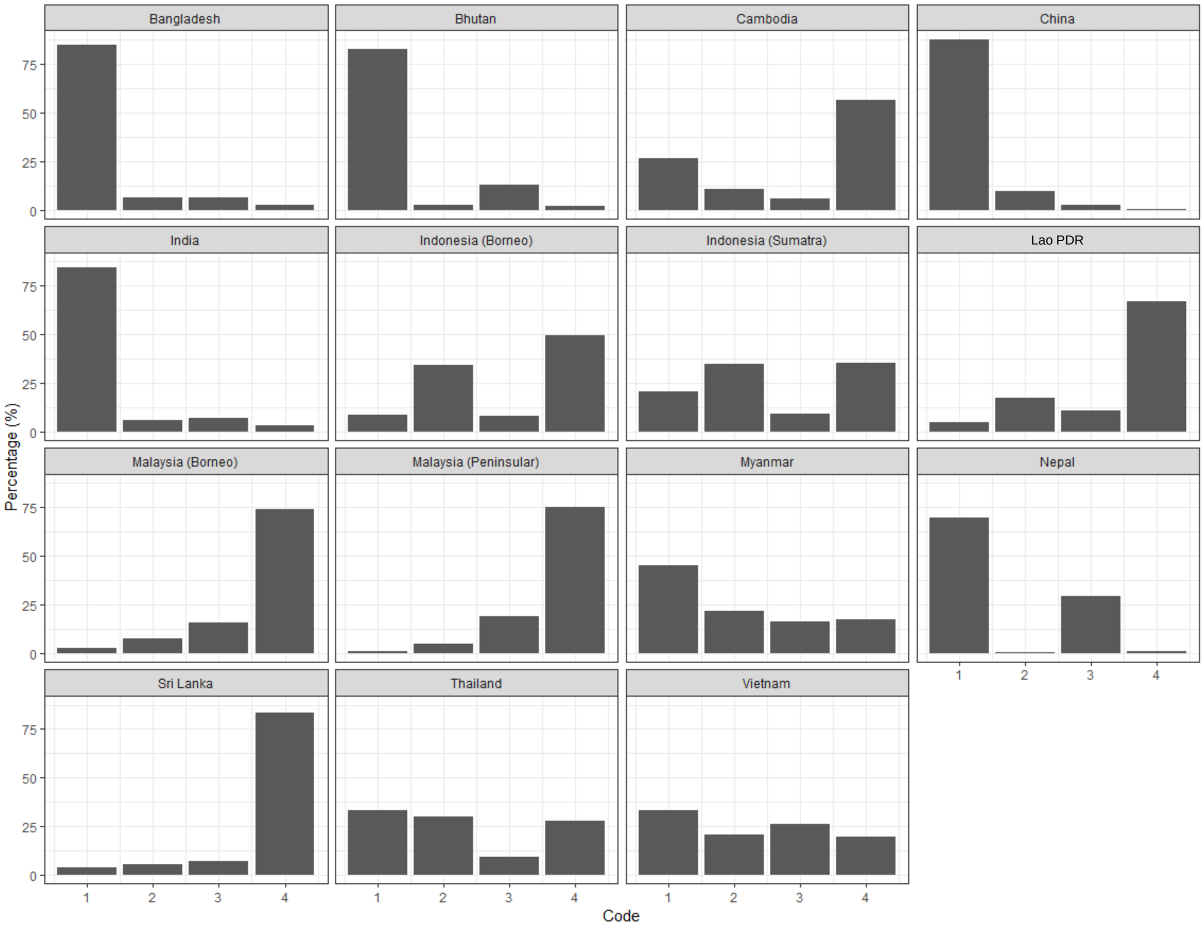

**References**

Calvin, K., Bond-Lamberty, B., Clarke, L., Edmonds, J., Eom, J., Hartin, C., Kim, S., Kyle, P., Link, R., Moss, R., McJeon, H., Patel, P., Smith, S., Waldhoff, S., & Wise, M. (2017). The SSP4: A world of deepening inequality. *Global Environmental Change*, *42*, 284–296. https://doi.org/10.1016/j.gloenvcha.2016.06.010

Hasegawa, T., Fujimori, S., Ito, A., Takahashi, K., & Masui, T. (2017). Global land-use allocation model linked to an integrated assessment model. *Science of The Total Environment*, *580*, 787–796. https://doi.org/10.1016/J.SCITOTENV.2016.12.025

Hurtt, G. C., Chini, L., Sahajpal, R., Frolking, S., Bodirsky, B. L., Calvin, K., Doelman, J., Fisk, J., Fujimori, S., Goldewijk, K. K., Hasegawa, T., Havlik, P., Heinimann, A., Humpender, F., Jungclaus, J., Kaplan, J., Krisztin, T., Lawrence, D., & Lawre, X. (2020). Harmonization of Global Land Use Change and Management for the Period 850-2100 (LUH2) for CMIP6. In *Earth System Grid Federation*.

O'Neill, B. C., Kriegler, E., Ebi, K. L., Kemp-Benedict, E., Riahi, K., Rothman, D. S., van Ruijven, B. J., van Vuuren, D. P., Birkmann, J., Kok, K., Levy, M., & Solecki, W. (2015). The roads ahead: Narratives for shared socioeconomic pathways describing world futures in the 21st century. *Global Environmental Change*, *42*, 169–180. https://doi.org/10.1016/j.gloenvcha.2015.01.004

O'Neill, B. C., Kriegler, E., Riahi, K., Ebi, K. L., Hallegatte, S., Carter, T. R., Mathur, R., & van Vuuren, D. P. (2014). A new scenario framework for climate change research: The concept of shared socioeconomic pathways. *Climatic Change*, *122*(3), 387–400. https://doi.org/10.1007/s10584-013-0905-2

Riahi, K., van Vuuren, D. P., Kriegler, E., Edmonds, J., O'Neill, B. C., Fujimori, S., Bauer, N., Calvin, K., Dellink, R., Fricko, O., Lutz, W., Popp, A., Cuaresma, J. C., KC, S., Leimbach, M., Jiang, L., Kram, T., Rao, S., Emmerling, J., … Tavoni, M. (2017). The Shared Socioeconomic Pathways and their energy, land use, and greenhouse gas emissions implications: An overview. *Global Environmental Change*, *42*, 153–168. https://doi.org/10.1016/j.gloenvcha.2016.05.009

Stehfest, E., Van Vuuren, D., Bouwman, L., & Kram, T. (2014). *Integrated assessment of global environmental change with IMAGE 3.0: Model description and policy applications.*

van Vuuren, D. P., Edmonds, J., Kainuma, M., Riahi, K., Thomson, A., Hibbard, K., Hurtt, G. C., Kram, T., Krey, V., Lamarque, J. F., Masui, T., Meinshausen, M., Nakicenovic, N., Smith, S. J., & Rose, S. K. (2011). The representative concentration pathways: An overview. *Climatic Change*, *109*(1), 5–31. https://doi.org/10.1007/S10584-011-0148-Z/TABLES/4

van Vuuren, D. P., Stehfest, E., den Elzen, M. G. J., Kram, T., van Vliet, J., Deetman, S., Isaac, M., Goldewijk, K. K., Hof, A., Beltran, A. M., Oostenrijk, R., & van Ruijven, B. (2011). RCP2.6: Exploring the possibility to keep global mean temperature increase below 2°C. *Climatic Change*, *109*(1), 95–116. https://doi.org/10.1007/S10584-011-0152-3/FIGURES/12

Wise, M., Calvin, K., Kyle, P., Luckow, P., & Edmonds, J. (2014). Economic and physical modeling of land use in GCAM 3.0 and an application to agricultural productivity, land, and terrestrial carbon. *Climate Change Economics*, *5*(2), 1450003. https://doi.org/10.1142/S2010007814500031
